# Monoterpene glucosides accumulated in *Eustoma grandiflorum* roots promote hyphal branching in arbuscular mycorrhizal fungi

**DOI:** 10.1101/2023.04.24.538035

**Authors:** Takaya Tominaga, Kotomi Ueno, Hikaru Saito, Mayumi Egusa, Katsushi Yamaguchi, Shuji Shigenobu, Hironori Kaminaka

**Affiliations:** Graduate School of Science and Technology, Nara Institute of Science and Technology, Ikoma, Nara 630-0192, Japan; Faculty of Agriculture, Tottori University, Tottori 680-8553, Japan; Functional Genomics Facility, NIBB Core Research Facilities, National Institute for Basic Biology, Okazaki, Aichi 444-8585, Japan; Unused Bioresource Utilization Center, Tottori University, Tottori, 680-8550, Japan

**Keywords:** arbuscular mycorrhizal symbiosis, *Eustoma grandiflorum*, gibberellin, hyphal branching, secoiridoid glucosides

## Abstract

Host plant-derived strigolactones trigger hyphal branching in arbuscular mycorrhizal (AM) fungi, initiating a symbiotic interaction between land plants and AM fungi. However, our previous studies revealed that gibberellin-treated *Eustoma grandiflorum* (Gentianaceae) activates rhizospheric hyphal branching in AM fungi using unidentified molecules other than strigolactones. In this study, we analyzed independent transcriptomic data of *E. grandiflorum* and found that the gentiopicroside (GPS) and swertiamarin (SWM), which are characteristic monoterpene glucosides in Gentianaceae, were highly biosynthesized in gibberellin-treated *E. grandiflorum* roots. Moreover, these metabolites considerably promoted hyphal branching in the Glomeraceae AM fungi *Rhizophagus irregularis* and *R. clarus*. GPS treatment also enhanced *R. irregularis* colonization of the monocotyledonous crop *Allium schoenoprasum*. Interestingly, these metabolites did not provoke the germination of the root parasitic plant *Orobanche minor*. Altogether, our study unveiled the crucial role of GPS and SWM in activating the symbiotic relationship between AM fungi and *E. grandiflorum*.

## Introduction

Various microbes reside in plants’ roots and influence their adaptation to environments, being either beneficial or detrimental to plants’ lifecycles (Bakker et al., 2018). For survival under such conditions, plants utilize secondary metabolites to control microbial communities and function in the narrow space around the roots, termed the rhizosphere. Recent studies revealed that several defense molecules, including saponins, are secreted into the rhizosphere of crop species, increasing the population of beneficial microbes (Fujimatsu et al., 2020; Nakayasu et al., 2021; Zhong et al., 2022). Regarding symbiotic microbes, arbuscular mycorrhizal (AM) fungi of the Glomeromycotina sub-phylum are the most general fungal partners of terrestrial plants (Brundrett and Tedersoo, 2018). AM fungi transfer inorganic phosphate from beyond the reach of root systems to host plants, thereby promoting plant growth (Luginbuehl and Oldroyd, 2017). Host plants are thus programmed to secrete strigolactones (SLs), which stimulate hyphal branching in AM fungi, in response to phosphate deficiency (Akiyama et al., 2005; Yoneyama et al., 2007). Furthermore, SL-deficient mutants fail to initiate and maintain AM symbiosis (Kretzschmar et al., 2012; Kobae et al., 2018; Kodama et al., 2022). Therefore, SLs have been considered representative signal molecules for establishing AM symbiosis in land plants.

Recently, it was clarified that plants properly regulate AM symbiosis via several phytohormones (Gutjahr, 2014). In model plants, gibberellin (GA) suppresses AM fungal colonization and the formation of highly branched hyphal structures for effective nutrient exchange, namely arbuscules, in a concentration-dependent manner (Takeda et al., 2015; Nouri et al., 2021). The repressor DELLA activating AM symbiosis, the degradation of which is triggered by GA, could be responsible for the inhibitory effects of GA (Davière and Achard, 2013; Foo et al., 2013; Pimprikar et al., 2016). However, our previous studies found that GA-treated *Eustoma grandiflorum* (Gentianaceae) roots promote the colonization of the model AM fungus *Rhizophagus irregularis* by increasing extraradical hyphal branching (Tominaga et al., 2020; Tominaga et al., 2021). Surprisingly, GA treatment significantly suppresses SL production in *E. grandiflorum* roots, as found in other model plants (Ito et al., 2017; Tominaga et al., 2021). Because *R. irregularis* exhibits no response to exogenous GA (Takeda et al., 2015; Tominaga et al., 2020), our findings indicate that other metabolites accumulated by or exuded from GA-treated *E. grandiflorum* roots activate *R. irregularis* branching.

In this study, we reanalyzed two independent transcriptomic datasets of *E. grandiflorum* roots (Tominaga et al., 2020; Tominaga et al., 2021) to identify novel branching-inducing factors other than SLs. The analysis revealed that GA-treated *E. grandiflorum* roots activated the production of gentiopicroside (GPS) and swertiamarin (SWM), characteristic monoterpenes called secoiridoid glucosides that exert antimicrobial and anti-inflammatory properties in Gentianaceae plants (Yu et al., 2004; Šiler et al., 2010). We next quantified the hyphal branching induced by standard GPS and SWM using three species of AM fungi. As a result, the Glomeraceae fungi *R. irregularis* and *R. clarus* displayed increased hyphal branching upon GPS and SWM exposure, whereas the Gigasporaceae fungus *Gigaspora margarita* did not respond to these agents. Interestingly, the colonization of *G. margarita* in *E. grandiflorum* roots was not promoted by GA treatment. We also found that exogenous GPS treatment significantly enhanced *R. irregularis* colonization in another crop species, namely *Allium schoenoprasum* (chive), without stimulating seed germination in a root parasitic plant. Therefore, our findings offer new insights into the role of well-known defense molecules produced by Gentianaceae plants when activating AM fungi.

## Results

### Colonization by two *Rhizophagus* fungi is promoted in GA-treated *E. grandiflorum* roots

Our recent works illustrated that the hyphal branching of *R. irregularis* is drastically promoted in GA-treated *E. grandiflorum* rhizospheres (Tominaga et al., 2020; Tominaga et al., 2021). Previous studies used *Gigaspora* fungi for branching assays because of their simpler hyphal structure compared to that of *R. irregularis* (Akiyama et al., 2005; Besserer et al., 2006; Tsuzuki et al., 2016). Hence, we first explored a suitable AM fungus for the assay to identify *E. grandiflorum*-derived branching factors. The extraradical hyphal branching, colonization, and hyphopodia formation of another *Rhizophagus* species, namely *R. clarus* was enhanced in GA-treated *E. grandiflorum* roots (Supplemental Fig. S1A). However, *G. margarita* displayed no positive responses to GA-treated *E. grandiflorum* roots (Supplemental Fig. S1B). The branch formation and colonization of both AM fungi were considerably suppressed in GA-treated *Lotus japonicus* roots (Supplemental Fig. S1, C and D). Because GA inhibited SL biosynthesis in *E. grandiflorum* and *L. japonicus* (Ito et al., 2017; Tominaga et al., 2021), these data support the existence of unknown molecules that stimulate *Rhizophagus* fungi in *E. grandiflorum*. In addition, it was predicted that *G. margarita* would not respond to *E. grandiflorum*-derived branching factors. Thereafter, we mainly used *R. irregularis* to quantify the hyphal branching-inducing activity of metabolites in this study.

### Secoiridoid glucoside biosynthesis is enhanced in GA-treated *E. grandiflorum* mycorrhizae

We reanalyzed two past independent RNA-seq datasets because GA-treated *E. grandiflorum* mycorrhizal roots accumulate secondary metabolites that stimulate hyphal branching in *Rhizophagus* fungi. This analysis indicated that the expression of 1513 genes was significantly increased in GA-treated *E. grandiflorum* mycorrhizae compared to the findings in the respective controls (Supplemental Table S1, Log_2_ Fold Change [Log_2_FC] > 1, false discovery rate [FDR] < 0.01). Gene Ontology (GO) enrichment analysis revealed increases in UDP-glucosyltransferase activity and the production of secologanin, a precursor of secoiridoid glucosides including GPS and SWM (Fig. 1A and Supplemental Table S2) (De Luca et al., 2014; Cao et al., 2016; Rai et al., 2016; Božunović et al., 2019). In addition, the GO terms included secologanin biosynthetic genes annotated as *7-deoxyloganetic acid glucosyltransferase* (*7DLGT*) and *secologanin synthase* (*SLS*) (De Luca et al., 2014) (Supplemental Table S3). Based on these data, we further analyzed the expression patterns of homologous secoiridoid biosynthetic genes in *E. grandiflorum*. A greater than 2-fold increase in expression (FDR < 0.05) was found for one *geranyl diphosphate synthase*, five *geraniol 8-hydroxylase*, one *8-hydroxygeraniol oxidoreductase*, five *7DLGT*, two *7-deoxyloganic acid hydroxylase*, and four *SLS* genes in *E. grandiflorum* mycorrhizae (Fig. 1B and Supplemental Table S3). Therefore, these results indicate that secoiridoid production is enhanced in GA-treated *E. grandiflorum* mycorrhizae.

**Figure 1.**
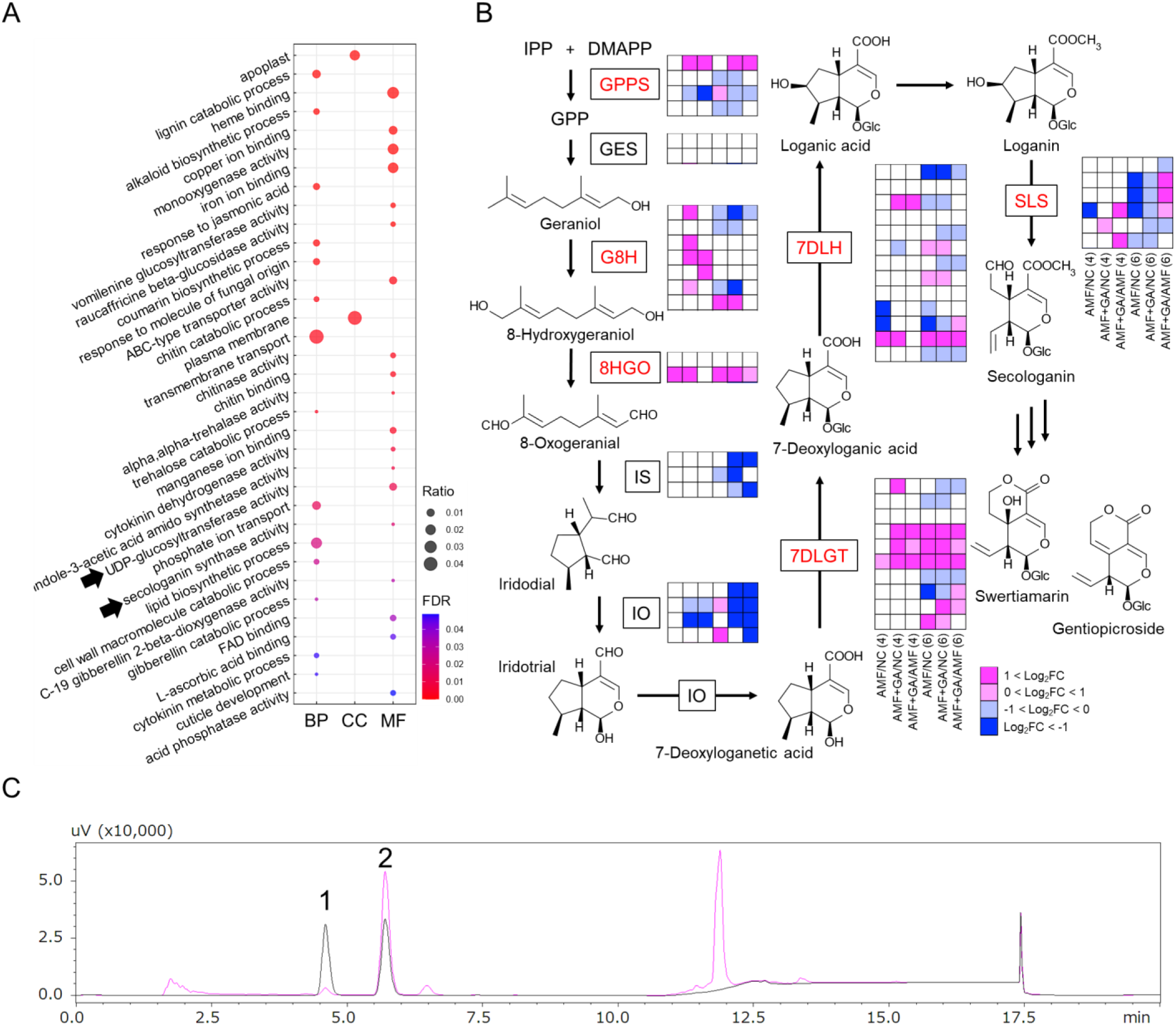
Transcriptional activation of the secoiridoid pathway in *Eustoma grandiflorum* upon GA treatment. A, GO enrichment analysis showing activated molecular function (MF), cellular component (CC), and biological process (BP) terms in GA-treated *E. grandiflorum* roots colonized by *Rhizophagus irregularis* at 4 and 6 weeks post-inoculation (wpi). Genes displaying significant expression (Log_2_FC > 1, FDR < 0.01) at either 4 or 6 wpi were analyzed. See also Supplemental Table S1 and S2. Black arrows indicate GO terms corresponding to secoiridoid biosynthesis. B, Expression pattern of genes involved in the secoiridoid biosynthetic pathway in *E. grandiflorum* (*n* = 3–4). NC, non-colonized roots; AMF, *R. irregularis* inoculation; AMF+GA, *R. irregularis* inoculation with GA treatment. Genes indicated in the boxes are involved in the secoiridoid pathway. Red letters represent genes upregulated by GA treatment. GPPS, geranyl diphosphate synthase; GES, geraniol synthase; G8H, geraniol 8-hydroxylase; 8HGO, 8-hydroxygeraniol oxidoreductase; IS, iridoid synthase; IO, iridoid oxidase; 7DLGT, 7-deoxyloganetic acid glucosyltransferase; 7DLH, 7-deoxyloganic acid hydroxylase; SLS, secologanin synthase. Magenta and blue denote positive and negative changes in the expression of each gene compared with NC or AMF, respectively (FDR < 0.05). C, Identification of SWM (peak 1; 4.6 min) and GPS (peak 2; 5.7 min) from the methanol extracts of 6-week-old axenic *E. grandiflorum* roots (magenta line) by HPLC. The black line indicates the peaks of the SWM and GPS standards. See also Supplemental Tables S1 and S2.

In Gentianaceae plants, secologanin is generally converted into GPS and SWM (Cao et al., 2016; Rai et al., 2016). However, these secoiridoid glucosides have not yet been detected in *E. grandiflorum*. Hence, we first investigated the accumulation of GPS and SWM in axenically grown *E. grandiflorum* roots. HPLC using a reverse-phase column detected two peaks from the methanol extracts of *E. grandiflorum* roots that matched the retention times of standard GPS and SWM (Fig. 1C). Moreover, peaks 1 and 2 had unique UV spectra that matched the SWM and GPS standard spectra (99.9% and 100%, respectively; Supplemental Fig. S2). These results illustrate that *E. grandiflorum* is capable of GPS and SWM biosynthesis.

### GA treatment increased GPS and SWM accumulation in *E. grandiflorum* roots

Because the ability of *E. grandiflorum* to biosynthesize GPS and SWM was confirmed, we subsequently quantified the effects of GA treatment on the levels of these compounds in *E. grandiflorum* roots. HPLC revealed that GPS content was significantly increased in *E. grandiflorum* mycorrhizae 4 weeks after GA treatment compared to that in mock-treated mycorrhizae (Fig. 2A, *P* = 0.0071). GA treatment also increased SWM accumulation in 4-week-old *E. grandiflorum* mycorrhizae compared to that in mock-treated roots (Fig. 2B, *P* = 0.052). These data suggest that exogenous GA promotes GPS and SWM biosynthesis in *E. grandiflorum* roots, consistent with the results of transcriptome analyses (Fig. 1 and Supplemental Table S2).

**Figure 2.**
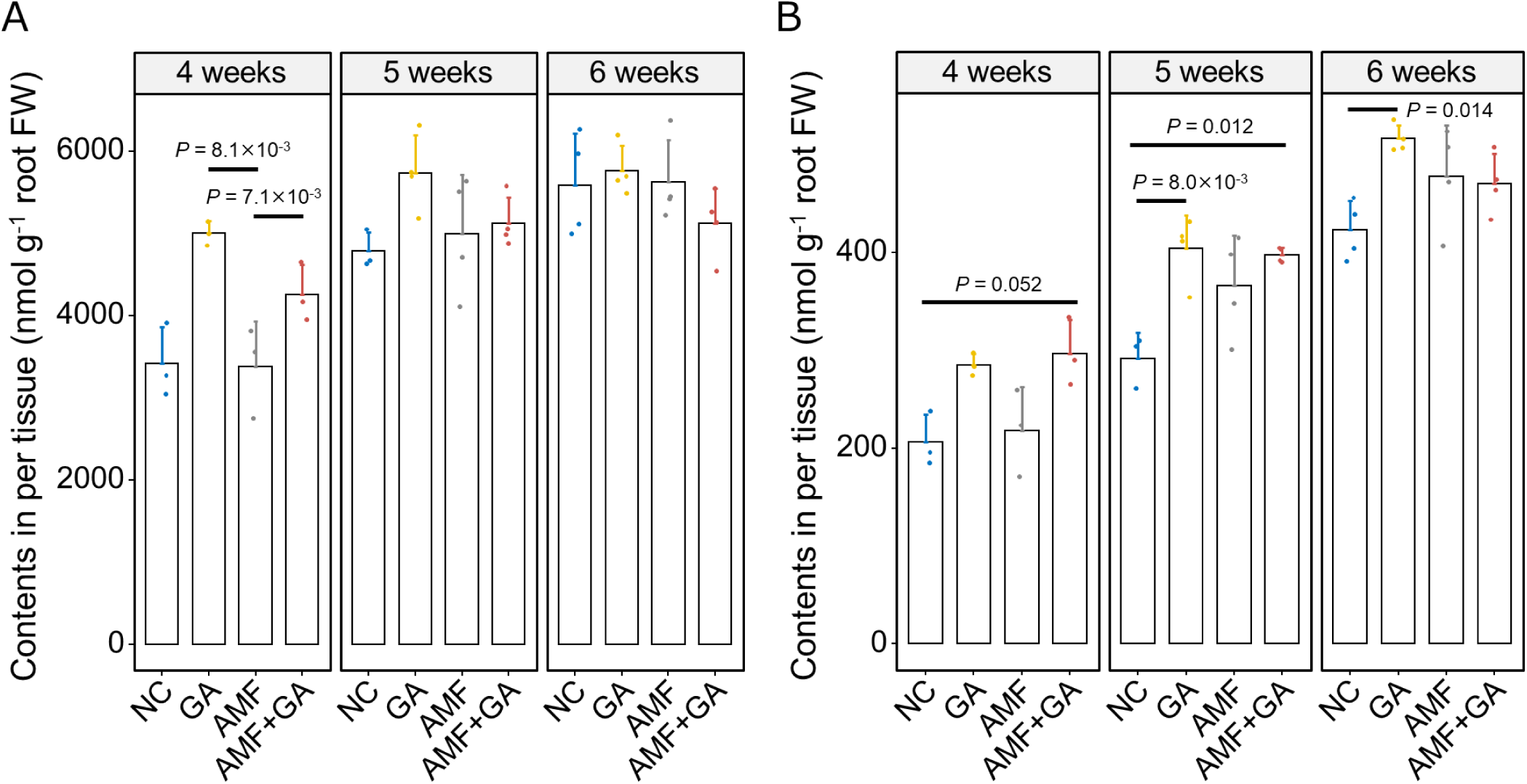
Effects of GA treatment on GPS and SWM content in *E. grandiflorum* roots. A and B, HPLC analysis of GPS (A) and SWM (B) extracted from *E. grandiflorum* roots at 4–6 weeks. The plants were treated with 0.01% ethanol for mock treatment and 1 µM GA_3_. NC, non-colonized roots; GA, GA treatment; AMF, *R. irregularis* inoculation; AMF+GA, *R. irregularis* inoculation with GA treatment. Bars indicate the means of GPS and SWM nmol (g root fresh weight [FW])^−1^, and error bars represent the standard deviation (*n* = 3–4 biologically independent samples). The significant differences among treatments were tested using Welch’s *t*-test with Bonferroni correction after confirming the normality of the data using the Shapiro–Wilk test.

The presence of highly branched *R. irregularis* hyphae in GA-treated *E. grandiflorum* rhizospheres (Tominaga et al., 2020) prompted us to hypothesize that *E. grandiflorum* roots secrete some signal compounds into soil. Thus, we quantified the levels of GPS and SWM in the root exudates; however, the levels of GPS and SWM in the root exudates were below the detection limits (GPS, 4.4 µM; SWM, 0.45 µM). Altogether, the branching factors promote hyphal branching in *R. irregularis* and *R. clarus* at lower concentrations than the detection limits.

### GPS and SWM increase hyphal branching in two *Rhizophagus* species

To investigate whether GPS and SWM can promote hyphal branching in *R. irregularis* and *R. clarus*, we applied an *in vitro* assay system as previously described (Kameoka et al., 2019). This assay enabled us to quantify the number of hyphal branches emerging from straight, elongating, thick hyphae (Fig. 3A). The number of *R. irregularis* branches was significantly increased by the synthetic SL *rac*-GR24 (GR24), in agreement with other studies (Cohen et al., 2013; Tsuzuki et al., 2016). Exogenous GPS and SWM treatment at concentrations of 1–100 nM promoted hyphal branching in *R. irregularis* (Fig. 3, B and C) despite their antifungal effects on the pathogenic fungus *Fusarium oxysporum* f. sp. *lycopersici* (Supplemental Fig. S3). The activities of GPS and SWM were comparable to that of GR24 *in vitro*. Moreover, these metabolites slightly increased the number of hyphal branches in *R. irregularis* at the femtomolar level (Supplemental Fig. S4, *P* > 0.085). Conversely, exogenous GA did not affect hyphal branching in the AM fungus at any examined concentration (Supplemental Fig. S5, *P* > 0.44) (Takeda et al., 2015; Tominaga et al., 2020).

**Figure 3.**
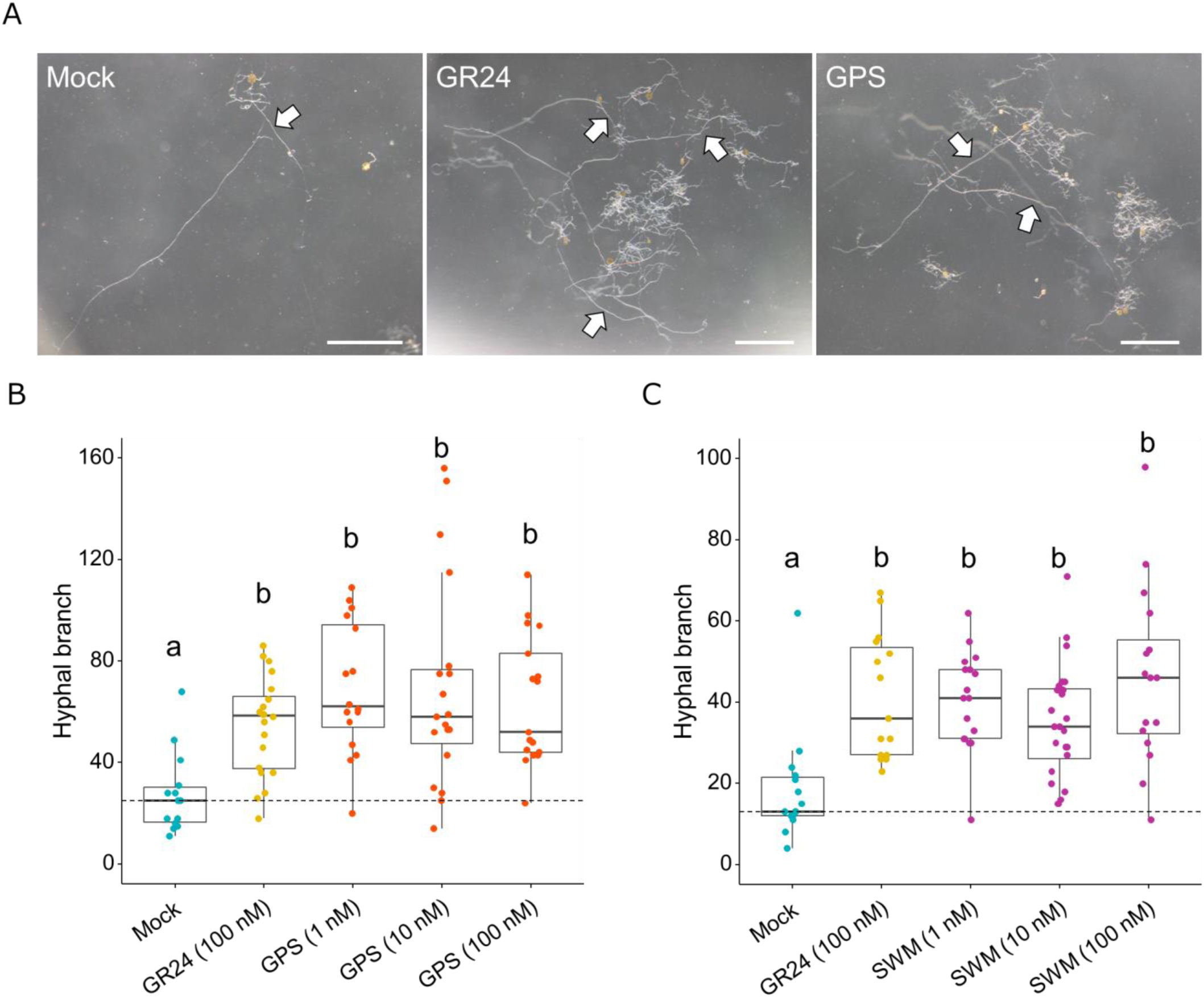
Quantification of hyphal branching-inducing activity using an in *vitro* assay. A, *R. irregularis* germinating spores treated with distilled water (Mock, left), 100 nM GR24 (middle), and 10 nM GPS (right) for 7 days. The hyphal branches on straight elongating thick hyphae (arrows) were counted. Scale bars, 1 mm. B and C, The number of *R. irregularis* hyphal branches in the presence of GPS (B) and WM (C). Data are shown as box plots with the 25th–75th percentiles (box), median (center line inside the box), and range (whiskers) [*n* = 14–20 (B) and *n* = 15–24 (C)]. Different letters indicate significant differences among treatments as determined by Wilcoxon’s rank-sum test with Bonferroni’s correction (*P* < 0.05).

To determine whether *R. irregularis* specifically responded to GPS or SWM, we further quantified the branching induced by other (seco)iridoid glucosides found in nature. The secologanin precursor loganin most strongly promoted hyphal branching in *R. irregularis*, and its effect was comparable to that of GR24 (Supplemental Fig. S6A, *P* < 0.0038). The hyphal branching-inducing activities of geniposide and oleuropein identified from *Gardenia jasminoides* (Wang et al., 2004) and *Olea europaea* (Soler-Rivas et al., 2000), respectively, were identical to that of mock treatment at all concentrations excluding oleuropein at 10 nM (Supplemental Fig. S6A, *P* = 0.0097). However, geraniol, an intermediate of these secoiridoid glucosides, did not change *R. irregularis* branching compared to the effects of mock treatment (Supplemental Fig. S6B). These data suggest that (seco)iridoid glucosides produced by Gentianaceae plants stimulate *R. irregularis* hyphal branching.

The different responses of *R. clarus* and *G. margarita* to GA-treated *E. grandiflorum* roots (Supplemental Fig. S1) suggested that the two AM fungi differentially respond to GPS and SWM. We thus treated *R. clarus* and *G. margarita* with GPS and SWM. *R. clarus* exhibited a significant but slight increase in hyphal branching in the presence of GPS and SWM (Supplemental Fig. S7A). By contrast, hyphal branching in *G. margarita* was not triggered by exogenous GPS and SWM at any concentration (Supplemental Fig. S7B). Interestingly, the response of *G. margarita* to GPS and SWM was consistent with our hypothesis that *G. margarita* is insensitive to branching factors derived from *E. grandiflorum* roots (Supplemental Fig. S1B). Therefore, GPS and SWM are representative compounds promoting the extraradical hyphal branching of *R. irregularis* and *R. clarus* in GA-treated *E. grandiflorum* roots.

### Transcriptome analysis of *R. irregularis* treated with GPS

To clarify the mechanisms underlying GPS-mediated hyphal branching in *R. irregularis*, we conducted RNA-seq analysis of GPS-treated *R. irregularis*. In this analysis, germinating *R. irregularis* spores were treated with 100 nM GR24 and GPS. The AM fungal hyphae highly branched, and the branches became entangled (Fig. 4A). Transcriptome analysis showed that 54 and 728 genes were transcriptionally upregulated (Log_2_FC > 1, FDR < 0.05) in *R. irregularis* by GPS and GR24 treatment, respectively (Fig. 4, B and C and Supplemental Table S4). The number of genes upregulated by GPS was smaller than that upregulated by GR24. Conversely, 96.3% of upregulated genes and 94.4% of downregulated genes were shared between GPS-treated and GR24-treated *R. irregularis* (Fig. 4C). These data indicate that *R. irregularis* exhibits partially similar transcriptional changes in response to GPS and GR24.

**Figure 4.**
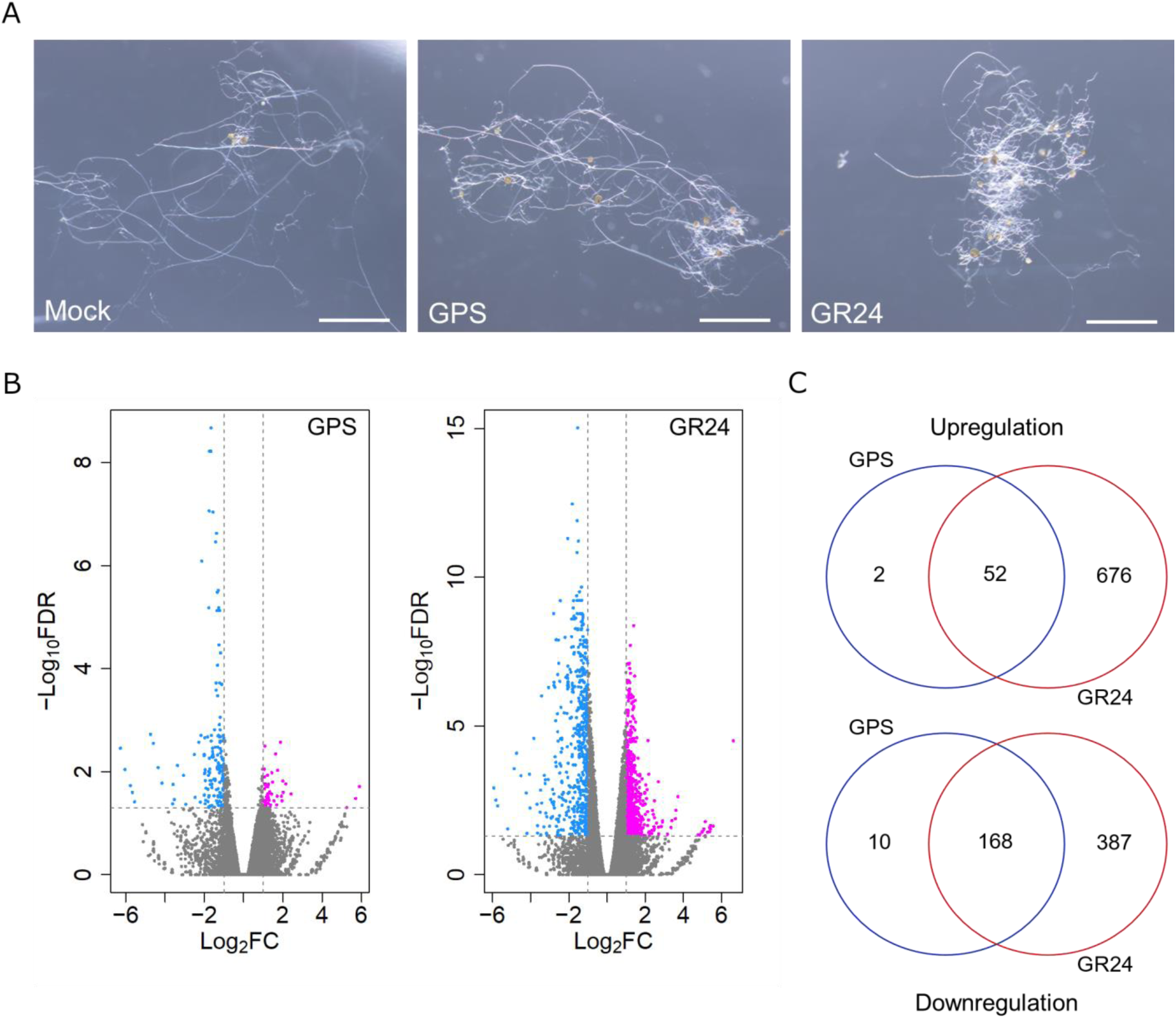
Transcriptional responses of GPS-treated *R. irregularis*. *R. irregularis* spores were germinated in M liquid medium for 5 days, followed by treatment with 100 nM GPS or GR24. After 8 days, the fungal RNA was extracted from 40,000 germinating spores. A, *R. irregularis* germinating spores and hyphae in each treatment. Scale bars, 1 mm. B, Volcano plots showing the distribution of the DEGs of *R. irregularis* treated with GPS (left) or GR24 (right). Horizontal lines represent that the FDR cut-off was set as 0.05, and vertical lines indicate that the Log_2_FC threshold was set as −1 and 1. The downregulated and upregulated DEGs are colored cyan and magenta, respectively. C, Venn diagrams displaying the expression patterns of the fungal DEGs upon GPS and GR24 treatment. Each treatment consisted of three biologically independent samples. See also Supplemental Table S4.

We further performed GO enrichment analysis to investigate the mechanism by which GPS triggers *R. irregularis* branching. As a result, 52 genes upregulated by both GPS and GR24 were highly enriched with cytoskeletal functions (Supplemental Table S5, *P* < 0.0092, FDR = 1). In addition, both treatments also activated protein serine/threonine kinase activity (GO: 0004674; Supplemental Table S5, *P* = 0.029, FDR = 1). These data reflect cytoskeletal function in the hyphal branching of filamentous fungi (Lichius et al., 2011). Contrarily, GPS and GR24 significantly downregulated 178 and 555 *R. irregularis* genes (Log_2_FC < 1, FDR < 0.05), respectively (Fig. 4, B and C and Supplemental Table S4). Many GO terms corresponding to respiration and mitochondrial activity were transcriptionally downregulated in response to GPS and GR24 (Supplemental Table S5, FDR < 0.05). These results contradicted previous findings showing the positive effects of GR24 on AM fungal mitochondrial activity (Besserer et al., 2006; Besserer et al., 2008). The AM fungal species and growth conditions applied in this study might be attributable to the negative impacts of GR24 on respiratory activity.

### GPS treatment enhances *R. irregularis* colonization in another crop species

GPS and SWM increased hyphal branching in *R. irregularis* and *R. clarus in vitro*; thus, we hypothesized that secoiridoid glucosides also have the ability to promote AM symbiosis in other host plants lacking GPS/SWM. Indeed, GPS treatment enhanced *R. irregularis* colonization in the monocotyledonous crop *A. schoenoprasum* (chive), which does not produce GPS/SWM (Fig. 5, A and B and Supplemental Fig. S8). GPS treatment did not affect the growth of chive seedlings and hyphal morphologies in chive roots (Fig. 5, A and B, *P* < 0.043).

**Figure 5.**
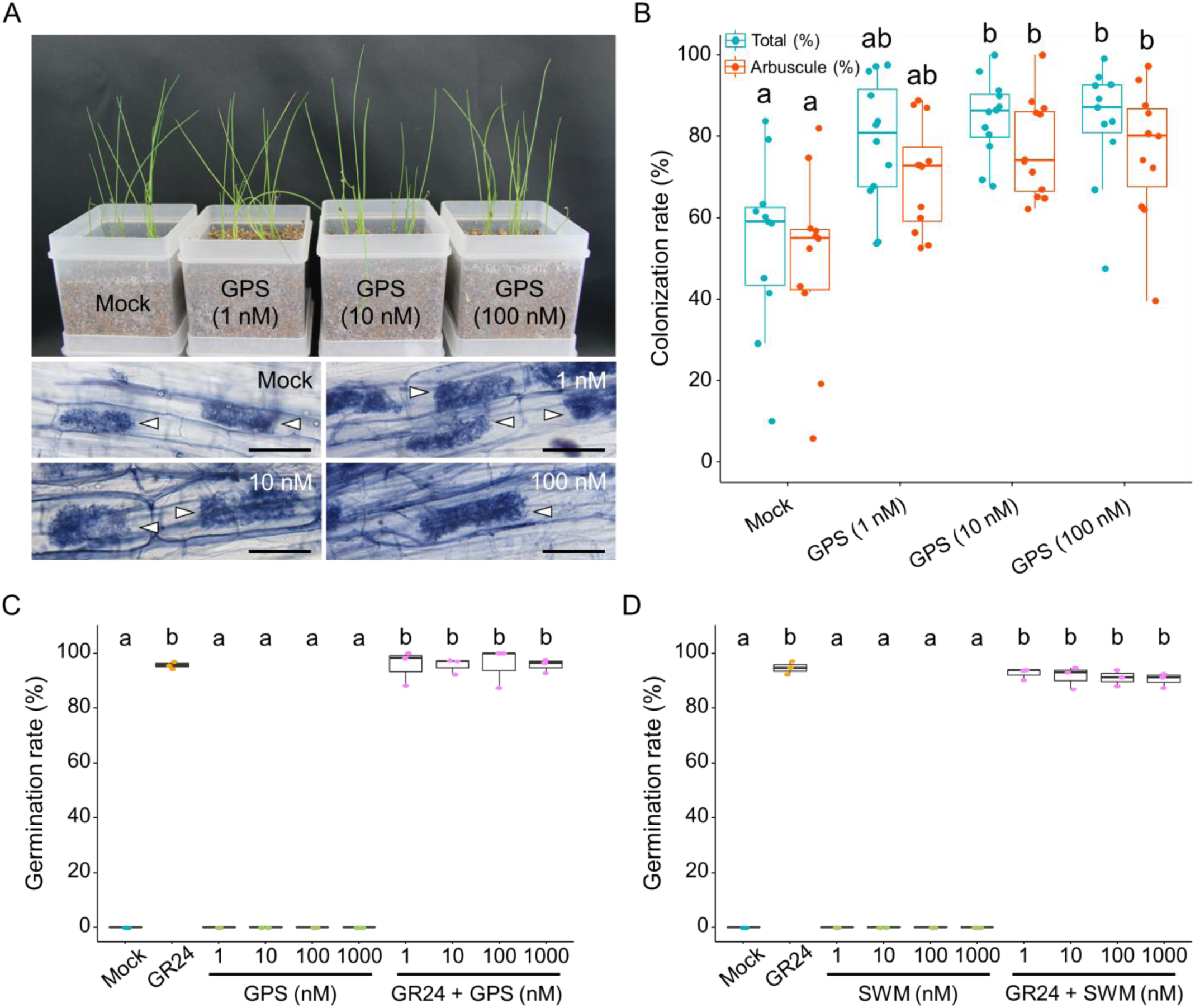
Exogenous GPS application improves *R. irregularis* colonization in chive roots without triggering *O. minor* seed germination. *A. schoenoprasum* (chive) roots inoculated with *R. irregularis* were harvested and observed after 1 month. A, Upper image shows the growth of chive seedlings treated with 0.01% ethanol and 1–100 nM GPS. The hyphal structures formed inside chive roots are displayed in the bottom pictures. Arrowheads indicate arbuscules. Scale bars, 50 µm. B, Colonization rates (%) of *R. irregularis* in chive roots. Green and orange plots present the total hyphal colonization and arbuscule formation rates, respectively. Significant differences among treatments as calculated using Wilcoxon’s rank-sum test with Bonferroni’s correction are indicated by different letters (*n* = 11–12, *P* < 0.05). C and D, Germination rate of *O. minor* seeds treated with 20 µL of distilled water (Mock), 1 µM GR24, and 1–1000 µM GPS (C) or SWM (D) (per disk) for 5 days (*n* = 3). *O. minor* seeds were also treated with 1 µM GR24 and 1–1000 nM GPS or SWM simultaneously. Data are shown as box plots with the 25–75th percentiles (box), the median (center line inside the box), and the minimum to maximum values (whiskers). Different alphabets indicate significant differences among treatments in Tukey test, *P* < 0.001.

Meanwhile, we applied a high-throughput bioassay to determine whether GPS could act as an SL using the root parasitic plant *Orobanche minor*, which germinates upon exogenous SL exposure (Ueno et al., 2014). Neither GPS nor SWM provoked *O. minor* germination (Fig. 5, C and D). These data suggest that GPS can activate the development of AM symbiosis without stimulating root parasitic plants.

## Discussion

Recently, it was reported that several defense molecules that taste bitter to humans maintain the healthy rhizosphere microbiota (Fujimatsu et al., 2020; Nakayasu et al., 2021; Zhong et al., 2022). Similarly, our findings revealed the positive effects of the antifungal and bitter secoiridoid glucosides GPS and SWM in *Rhizophagus* AM fungi (Kumarasamy et al., 2003; Šiler et al., 2010). When hydrolyzed by *β*-glucosidase, which is widely conserved among plants, fungi, and insects (Ketudat Cairns and Esen, 2010), secoiridoid glucosides are readily converted into toxic aglycones covalently binding to nucleotides and proteins (Konno et al., 1999; Kim et al., 2000; Dobler et al., 2011). Furthermore, cytosolic plant *β*-glucosidase is thought to hydrolyze secoiridoid glucosides when plant cells are damaged (Dobler et al., 2011). However, obligate biotrophic AM fungi have lost some polysaccharide hydrolases, including *β*-glucosidase (Tisserant et al., 2013; Kobayashi et al., 2018). Together with the non-destructive infection of AM fungi (Genre et al., 2008), secoiridoid glucosides accumulated in *E. grandiflorum* roots would be stable and non-toxic during AM symbiosis. These findings propose the bidirectional functions of GPS and SWM, namely the deterrence of pathogens and reinforcement of the symbiotic interaction (Fig. 6). Meanwhile, this study revealed no response of *G. margarita* to GPS/SWM. Because Gigasporaceae (genus *Gigaspora*) fungi sometimes depress plant growth and demand more carbon than Glomeraceae (genus *Rhizophagus*) fungi (Buwalda and Goh, 1982; Lendenmann et al., 2011; Kaur et al., 2022), GPS and SWM might contribute to attracting more cooperative AM fungi. This hypothesis should be tested in a wide range of AM fungal species because our current study only used three species of AM fungi.

**Figure 6.**
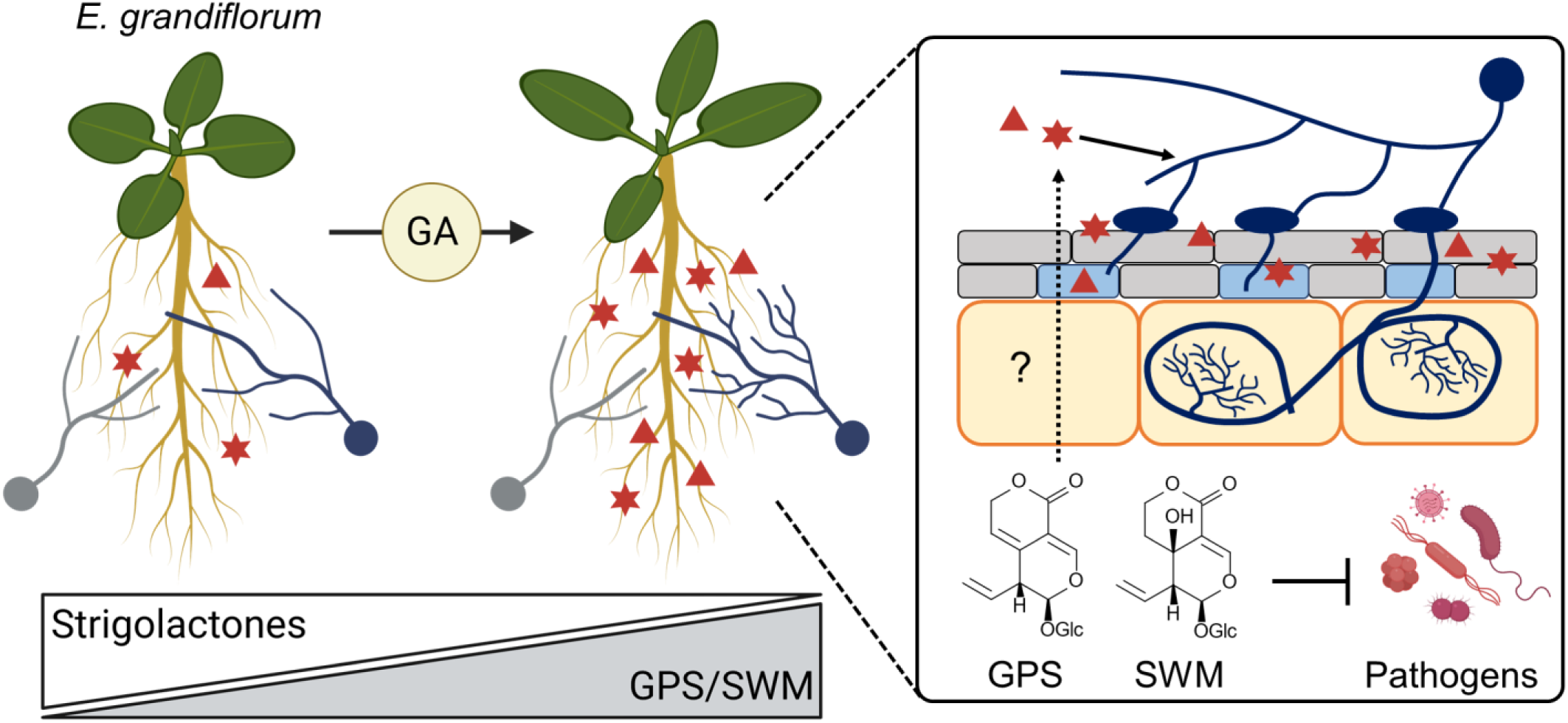
Graphical summary of the roles of GPS and SWM in AM symbiosis in *E. grandiflorum*. *E. grandiflorum* would not need SLs to associate with AM fungi in the presence of GA. Moreover, *R. irregularis* and *R. clarus* (blue) highly branch around GA-treated *E. grandiflorum* roots, unlike the phylogenetically distant AM fungus *G. margarita* (gray). GA activates the biosynthesis of monoterpenes GPS (red stars) and SWM (red triangles) in *E. grandiflorum* roots. These antimicrobial metabolites promote branch formation in two *Rhizophagus* fungi but do not alter *G. margarita* hyphal branching, consistent with the responses to GA-treated *E. grandiflorum* roots. Because the secretion of GPS/SWM has not been confirmed, their transport is shown as a dotted line. The light blue-colored cells in the right image represent hypodermal passage cells in which AM fungal hyphae constantly penetrate before colonizing the root cortex. This figure was created using BioRender.com.

Canonical SLs are terpenoid lactones featuring a tricyclic ABC lactone and a methyl butenolide connected by an enol ether (Yoneyama et al., 2018). Interestingly, GPS and SWM also possess one lactone ring. Moreover, it has been revealed that the lactone-forming coumarin scopoletin significantly stimulates *R. irregularis* hyphal elongation and metabolic activity in a concentration-dependent manner (Cosme et al., 2021). By contrast, lactone-lacking geraniol and other secoiridoid glucosides displayed no or weak hyphal branching-inducing activity excluding for the iridoid glucoside loganin (Supplemental Fig. S6). These findings imply that *Rhizophagus* fungi respond to the lactone ring of GPS and SWM. However, the lactone ring in the C-ring of SLs is dispensable for hyphal branching induction in *R. irregularis* (Cohen et al., 2013). In addition, GPS and SWM do not have the methyl butenolide and enol ether bridge required for root parasitic plant germination (de Saint Germain et al., 2013). This would explain why *O. minor* seeds did not respond to GPS (Fig. 5, C and D). Further studies are needed to determine whether the mechanisms underlying the recognition of SLs and GPS/SWM by *Rhizophagus* fungi are common.

More than 90% of positively expressed genes upon GPS treatment were also upregulated by GR24, and these genes were enriched in GO terms corresponding to cytoskeletal function and kinase activity (Fig. 4C and Supplemental Table S5). These functions are known to be activated during the host recognition of *R. irregularis* (Nadal et al., 2017; Tominaga et al., 2021). This study also confirmed the negative effects of GPS and GR24 on mitochondrial respiration activity in *R. irregularis* despite the positive impact of GR24 on AM fungal mitochondrial biogenesis (Besserer et al., 2006; Besserer et al., 2008). Our transcriptome analysis might have masked the responses of *R. irregularis* hyphae to GPS and GR24 because *R. irregularis* spores containing numerous nuclei feature distinct transcriptomes from hyphae (Kameoka et al., 2019). In addition, this study could not confirm the secretion of GPS or SWM from *E. grandiflorum* roots to the rhizosphere. The slight hyphal branching-inducing activity of GPS and SWM in the femtomolar range (Supplemental Fig. S3) suggests that their levels in *E. grandiflorum* root exudates are also too low to be detected. However, *Catharanthus roseus* roots secrete monoterpene indole alkaloids produced via the secoiridoid pathway (Nakabayashi et al., 2021), implying the possibility of GPS/SWM secretion from *E. grandiflorum* roots.

In conclusion, our data revealed that the representative monoterpenes of *E. grandiflorum*, namely GPS, and SWM, are key metabolites promoting *Rhizophagus* fungal branching. This finding further provides knowledge of the bidirectional functions of defense molecules in stimulating symbiotic partners. GPS-promoted AM fungal colonization in chive roots suggests their utilization as biostimulants that do not provoke Orobanchaceae parasitic plant germination (Fig. 5). On the contrary, the observation of successful AM symbiosis in GA-treated *E. grandiflorum* implies that the plant has evolved to undergo AM symbiosis when GA signaling is activated, such as that occurring in shaded areas (Yang and Li, 2017). Shady conditions also suppress SL biosynthesis, failing to accommodate AM fungi effectively (Nagata et al., 2015; Ge et al., 2022). Therefore, investigating whether GPS and SWM are genuinely involved in AM symbiosis under activated GA signaling would be interesting, considering the tandem duplication of secoiridoid biosynthetic genes in the Gentianales order and Gentianaceae family (Rai et al., 2021; Li et al., 2022; Zhou et al., 2022).

## Materials and methods

### Chemicals

GPS (>97.0%), SWM (>98.0%), loganin (>98.0%), geniposide (>95.0%), and oleuropein (>98.0%) were purchased from Tokyo Chemical Industry Co. (Tokyo, Japan). Methanol (HPLC grade, ≥99.7%), acetone (reagent grade, 99.5% purity), geraniol (>97.0%), and GA_3_ (>85.0%) were obtained from FUJIFILM Wako Pure Chemical Corp. (Osaka, Japan). GPS and GA_3_ dissolved in ethanol (reagent grade, 99.5% purity) were used to treat the examined plants by diluting them in 1/10 Hoagland solution at the indicated concentrations. In addition, we used the synthetic SL *rac*-GR24 (GR24) (>98.0%), which was synthesized by StrigoLab (Torino, Italy).

### Growth conditions of plant and fungal materials

The seeds of *L. japonicus* “Miyakojima” MG-20, *A. schoenoprasum* (chive), and *E. grandiflorum* cv. Pink Thumb were sterilized and germinated as described in our previous reports (Tominaga et al., 2020). The examined host plants were transplanted to boxes containing 300 mL of autoclaved mixed soil (river sand/vermiculite, 1:1). The plants were grown in a growth chamber under 14 h light/10 h dark cycles at 25°C for 4– 6 weeks in the presence of 1/10 Hoagland solution containing 100 µM NH_4_H_2_PO_4_. Chive seedlings were cultured with 1/5 Hoagland solution (20 µM NH_4_H_2_PO_4_) when we investigated the effects of GPS on AM symbiosis. GA_3_ and GPS diluted in ethanol were added to the Hoagland solutions at the indicated concentrations.

We inoculated the examined plants with *R. irregularis* DAOM197198 (Premier Tech, Quebec, Canada) by mixing 1000 spores in the soil mixture. Concerning the other AM fungal species, 50 spores of *R. clarus* and 15 spores of *G. margarita* were directly inoculated onto the host roots. *R. clarus* HR1 (MAFF520076) and *G. margarita* K-1 (MAFF520052) were obtained from the Genebank Project, National Agriculture and Food Research Organization of Japan. *R. clarus* and *G. margarita* were cultivated with *Medicago sativa* L. and *Trifolium pratense*, respectively. The soil inoculants were dried after 3 months and stored at 4°C until use. *R. clarus* and *G. margarita* spores were collected through 106- and 250-µm pore size sieves, respectively. Before use, the spores were sterilized with 1% (v/v) NaClO and 0.04% (v/v) Tween-20 for 20 min. As described previously (Tominaga et al., 2020), we evaluated AM fungal colonization rates (%) by staining harvested roots with 0.05% trypan blue diluted in lactic acid.

### Quantification of the bioactivity of chemicals

To quantify the hyphal branching-inducing activity of compounds against *Rhizophagus* fungi, we used a previously described method (Kameoka et al., 2019) with some modifications. Hyphae fragments in spore suspensions were removed via centrifugation in Gastrografin (Bayer Yakuhin, Osaka, Japan) solution before use (Furlan et al., 1980). Approximately eight spores of *R. irregularis* or *R. clarus* were incubated for 5 min on 350 μL of 0.4% (w/v) Phytagel (Sigma-Aldrich, St Louis, MO, USA) containing M medium (Hildebrandt et al., 2002) in a 24-well plate. Each aliquot of M medium was gently covered with 150 µL of liquid 0.3% (w/v) Phytagel in 3 mM MgSO_4_·7H_2_O cooled at 40°C. The AM fungal spores were germinated at 25°C in the dark for 5 days. We prepared at least three wells for each treatment in this study.

GPS, SWM, and three other secoiridoid glucosides diluted in distilled water were filtered through 0.45-µm PTFE filters (Shimadzu Co., Kyoto, Japan). Immediately, 200 µL of the axenic solutions were directly poured onto the gels containing the germinated AM fungal spores. We treated AM fungal spores with sterilized distilled water and 100 nM GR24 as a mock treatment. Then, 10 µM GR24 dissolved in acetone was dried in a SpeedVac DNA130 vacuum concentrator (Thermo Fisher Scientific, Waltham, MA, USA) at 35°C for 5 min. After removing acetone, GR24 was immediately redissolved at 100 nM in distilled water and sterilized as previously described. AM fungi treated with the chemicals were incubated at 25°C in the dark for 7–10 days. The number of hyphal branches emerging from straight elongating thick hyphae was counted under an SZX16 stereomicroscope (Olympus, Tokyo, Japan).

A single *G. margarita* spore was germinated on 0.2% (w/v) Phytagel in 3 mM MgSO_4_·7H_2_O at 30°C in the dark for 5–7 days. GPS and SWM dissolved in methanol were loaded onto 6-mm glass fiber disks at 0.1–10 µg/disk. After the solvent was dried entirely, the disks were placed near *G. margarita* hyphae as previously described^8^. The hyphae treated with the examined chemicals were microscopically observed. The root exudates collected from approximately 90 *T. pratens*e plants were used as a positive control because GR24 has lower ability to induce hyphal branching in *G. margarita* than other natural SLs (Akiyama et al., 2010). *T. pratense* seedlings were hydroponically grown with tap water for 20 days. The tap water medium (750 mL) was partitioned three times against an equal volume of ethyl acetate. The ethyl acetate extracts were dried over anhydrous Na_2_SO_4_, redissolved in acetone, and stored at 4°C until use.

### Germination assay of pathogenic fungal bud cells and root parasitic plant seeds

*F. oxysporum* f. sp. *lycopersici* strain JCM12575 bud cells were prepared as previously described (Egusa et al., 2019). *F. oxysporum* bud cells were treated with 1/2 potato dextrose broth containing 1–1000 nM GPS or SWM and incubated at 25°C in the dark for 12 h. The germinated bud cells were counted under a BX53 light microscope (Olympus). A germination assay of *Orobanche minor* seeds was conducted following our previous study (Tominaga et al., 2021). For simultaneous treatment with GR24 and GPS, 20 µL of 1 µM GR24 were added onto a 6-mm glass fiber disk, followed by the addition of 20 µL of GPS stock diluted at the indicated concentrations.

### Quantification of secoiridoid glucosides in *E. grandiflorum* roots

Fresh *E. grandiflorum* roots collected from two seedlings were weighted. The harvested roots were homogenized in a clean tube (INA-OPTIKA, Osaka, Japan) containing two beads and an aliquot of methanol using ShakeMan6 (Bio-Medical Science, Tokyo, Japan). The concentration of each sample was normalized by adding methanol at 50 mg root fresh weight (FW) mL^−1^. After extracting the root contents at room temperature overnight, the slurries were centrifuged at 13,000 rpm for 5 min. The supernatants were collected and filtered through 0.45-µm PTFE filters (Shimadzu). The endogenous levels of GPS and SWM were analyzed on an LC-2030C HPLC system (Shimadzu) equipped with a COSMOSIL 5C_18_-MS-II Packed Column (4.6 × 150 mm, 5 µm particle size; Nacalai Tesque, Kyoto, Japan) and a COSMOSIL Guard Column 5C_18_-MS-II (4.6 mm × 10 mm, 5 µm particle size; Nacalai Tesque) at 30°C. The mobile phases were Milli-Q water (solvent A) and methanol (solvent B), and the elution program was 30% B from 0 to 8 min, 30%–100% B (linear gradient) from 8 to 10 min, 100% B from 10 to 15 min, and 30% B from 15.01 to 20 min. The flow rate was 1 mL min^−1^, and the detection wavelength was 254 nm.

### Preparation of root exudates from *E. grandiflorum*

Four-week-old *E. grandiflorum* seedlings were transplanted into boxes containing 300 mL of washed and autoclaved river sand (0–1 mm particle size). *R. irregularis* spores and GA_3_ were mixed as previously mentioned. Distilled water (50 mL) was poured onto the river sand, and the filtrate was collected using a vacuum pump 4 and 6 weeks after transplanting. The collected samples were centrifuged at 13,000 rpm for 10 min and filtered with 0.45-µm PTFE membranes. The clear filtrates were loaded onto a Sep-Pak C18 Plus Short cartridge (Waters, Milford, MA, USA) and washed with 20 mL of distilled water. Metabolites were extracted with 6 mL of methanol from the sorbent. The extracted samples were stored at 4°C until use. Finally, the metabolites were dried and redissolved in distilled water at 50 mg root FW mL^−1^. The samples were immediately passed through 0.45-µm PTFE filters and used for the bioassay.

### RNA extraction from AM fungal spores and hyphae and RNA-seq

Five thousand *R. irregularis* spores were inoculated in 2.4 mL of M liquid medium in each well of a six-well plate and incubated at 25°C for 5 days in the dark. GR24 or GPS was added to the germinating spores at a final concentration of 100 nM. *R. irregularis* spores and hyphae in eight wells (40,000 spores per sample) were collected on a cell strainer after 8 days. The fungal sample was immediately frozen in an RNase-free tube containing two beads in liquid nitrogen. The frozen spores and hyphae were homogenized in ShakeMan6. RNA extraction and genomic DNA removal were performed using a Total RNA Extraction Kit (Plant) (RBC Bioscience, New Taipei, Taiwan) and DNase I (Takara Bio, Shiga, Japan) following the manufacturers’ protocol. The quality and quantity of the total RNA were checked using a Qubit RNA HS Assay Kit and Qubit 2.0 Fluorometer (Thermo Fisher Scientific) before sequencing. An RNA-seq library was constructed from the total RNA using an MGIEasy RNA Directional Library Prep Set (MGI, Shenzhen, China). RNA-seq with strand-specific and paired-end reads (150 bp) was performed using DNBSEQ-T7RS by Genome-Lead Co. (Takamatsu, Kagawa, Japan).

### Transcriptome analyses

The raw reads obtained from 4- and 6-week-old *E. grandiflorum* roots (Tominaga et al., 2020; Tominaga et al., 2020; Tominaga et al., 2021) and *R. irregularis* (this study) were filtered using Fastp v0.23.2 (Chen et al., 2018) to remove low-quality reads and adapter sequences. The filtered reads were mapped to the assembled *E. grandiflorum* 10B-620 (Shirasawa et al., 2023) and *R. irregularis* genome data (Maeda et al., 2018) using STAR v2.6.1d (Dobin et al., 2013) (Supplemental Table S6). Afterward, we quantified the number of reads aligned to the reference genome using featureCounts v2.0.1 (Liao et al., 2014). EdgeR v3.38.1 (Robinson et al., 2010) statistically calculated the fold change (FC) in gene expression and FDR. Differentially expressed genes (DEGs) were sorted using a Venn diagram (http://bioinformatics.psb.ugent.be/webtools/Venn). The GO pathways of each *E. grandiflorum* and *R. irregularis* gene were annotated using Blast2GO v6.0.3 (Conesa et al., 2005) to perform enrichment analysis. We first obtained a non-redundant (nr) ncbi-blast-dbs database from NCBI (https://github.com/yeban/ncbi-blast-dbs.git). After that, a Blastp search against the nr database was performed by DIAMOND v0.9.14 (Buchfink et al., 2021) using *E. grandiflorum* and *R. irregularis* protein sequences as query data. The resulting file was subjected to the Blast2GO program, and GO pathways were annotated with the default setting.

### Identification and analysis of iridoid biosynthesis genes in *E. grandiflorum*

A local tBlastx search (ncbi-blast-2.11.0, https://blast.ncbi.nlm.nih.gov/Blast.cgi) against *E. grandiflorum* genomic sequence using nucleotide sequences obtained from the Gentianales model plant *C. roseus* identified several iridoid biosynthesis genes with an E-value cut-off of 1e−100 (Supplemental Table 3). Only several genes with the same annotation as the queries were selected for the subsequent analysis.

### Quantification and statistical analysis

To quantify AM fungal colonization in host roots, we considered 10 pieces of approximately 10-mm root fragments collected from one seedling as one biological replicate (*n*, indicated in the figure legends). For RNA-seq of *R. irregularis*, we treated one pool of total RNA extracted from 40,000 spores as one biological replicate and prepared three biological replicates for each treatment. This study considered *R. irregularis* genes that fulfilled |Log_2_FC| ≥ 1 and FDR < 0.05 as DEGs. When we conducted the germination assay of *O. minor*, one glass filter disk with *O. minor* seeds was considered one biological replicate. The *O. minor* germination rate is shown as the mean of three biological replicates. All statistical analyses were conducted using R software v4.2.0. The differences in hyphal branching induced among the treatments were examined using Wilcoxon’s rank-sum test regarding the number of hyphal branches and root colonization rates. *P* values were corrected by the Bonferroni method for multiple comparisons. The Shapiro–Wilk test was used to examine the normality of GPS and SWM concentrations in *E. grandiflorum* roots, and the differences were tested using Welch’s *t*-test corrected by the Bonferroni method. The differences in the germination rates of *O. minor* and *F. oxysporum* f. sp. *lycopersici* were checked using Tukey’s test.

### Data availability

The original contributions presented in this study were publicly available. The RNA sequence data obtained from 4-week-old *E. grandiflorum* roots have been deposited into the DDBJ Sequence Read Archive under the accession numbers DRA010085 and DRA015766. The sequence of 6-week-old *E. grandiflorum* roots have previously been submitted to the DDBJ (DRA012117)(Tominaga et al., 2021). The RNA sequence data obtained from *R. irregularis* have also been available in the DDBJ (DRA015767).

## Acknowledgments

We would like to thank the National BioResource Project (Legume Base), Dr. Tsutomu Arie (Tokyo University of Agriculture and Technology), and Dr. Satoko Yoshida (Nara Institute of Science and Technology) for kindly providing *L. japonicus* seeds, *F. oxysporum* f. sp. *lycopersici*, and *O. minor* seeds, respectively. We thank Dr. Akifumi Sugiyama (Kyoto University), Dr. Shun Sakuma (Tottori University), and Dr. Hiromu Kameoka (CAS Center for Excellence in Molecular Plant Science) for critically reading our manuscript and giving valuable comments. This work was supported by the NIBB Cooperative Research Programs (Next-generation DNA Sequencing Initiative: 21-301, 22NIBB403), by the JST Adaptable and Seamless Technology transfer program through Target driven R&D (A-STEP) (Grant No. JPMJTM22DQ to H.K.), and a JSPS KAKENHI Grant-in-Aid for JSPS Fellows (Grant No. 20J21994 to T.T.). The graphical summary was created using BioRender.com.

## Author contributions

T.T. and H.K. designed the research; T.T. and H.S. performed the experiments, assisted by K.U. contributed to HPLC analyses and by M.E. contributed to antifungal activity assay; T.T., K.Y., and S.S. designed and performed the bioinformatic analyses; T.T and H.K. wrote the manuscript with comments from all authors.

## Supplementary data

**Supplemental Figure S1.** Different effects of GA treatment on AM symbioses between *E. grandiflorum* and *L. japonicus*. *E. grandiflorum* and *L. japonicus* roots inoculated with *R. clarus* or *G. margarita* were observed at 5 weeks post-inoculation. The plants were treated with 0.01% ethanol (Mock) or 1 µM GA_3_ (GA). A and B, *E. grandiflorum* roots were colonized by *R. clarus* (A) and *G. margarita* (B). Arrows denote extraradical hyphae adhering to *E. grandiflorum* roots. Scale bars, 5 mm (A) and 1 mm (B). Below graphs show the colonization rates (%) and hyphopodia number (mm^−1^) of *R. clarus* (A) and *G. margarita* (B) infecting *E. grandiflorum*. C and D, Colonization rates (%) and hyphopodia number (mm^−1^) of *R. clarus* (C) and *G. margarita* (D) infecting *L. japonicus*. Green and orange plots present the total hyphal colonization and arbuscule formation rates, respectively. Data are shown as box plots with the 25th –75th percentiles (box), median (center line inside the box), and range (whiskers). Asterisks indicate significant differences in GA-treated plants compared to mock-treated plant as determined using Wilcoxon’s rank-sum test (**: *P* < 0.01, *n* = 5–6).

**Supplemental Figure 2.** HPLC analyses of peaks 1 and 2 in Fig. 1C. A and B, Magenta lines represent the UV spectra of peaks 1 (A) and 2 (B) from the methanol extracts of 6-week-old axenic *E. grandiflorum* roots (Fig. 1C). Black lines indicate the UV spectra of SW (A) and GPS (B) standards. UV spectra of peaks 1 and 2 matched the SWM and GPS standard spectra (99.9% and 100%), respectively.

**Supplemental Figure 3.** Antifungal activity of GPS and SWM against *F. oxysporum*. A, Images of *F. oxysporum* f. sp. *lycopersici* bud cells treated with distilled water (left) or 10 nM GPS (right). Arrows indicate germinating bud cells, and arrowheads denote bud cells that did not germinate by 12 h. Scale bars, 100 µm. B and C, Germination rates of *F. oxysporum* bud cells treated with 1–1000 nM GPS (B) or SWM (C). Data are shown as box plots with the 25th–75th percentiles (box), the (center line inside the box), and range (whiskers). Different letters indicate significant differences among treatments as determined using the Tukey test (*P* < 0.001, *n* = 3).

**Supplemental Figure 4.** Hyphal branching induced by GPS and SWM in the femtomolar range. A and B, Images showing *R. irregularis* hyphae treated with femtomolar level GPS (A) and SWM (B). Distilled water was used as a mock control. Scale bars, 1 mm. C and D, The number of hyphal branches of *R. irregularis* treated with GPS (C) and SWM (D). There was no difference among treatments as determined using Wilcoxon’s rank-sum test with Bonferroni’s correction (*n* = 5–11). Data are shown as box plots with the 25th–75th percentiles (box), median (center line inside the box), and range (whiskers).

**Supplemental Figure 5.** Hyphal branching-inducing activity of GA. The effect of exogenous GA_3_ on *R. irregularis* hyphal branching. For the mock treatment, 0.01% ethanol was supplied to the fungus (*n* = 9–13). Different letters indicate significant differences among treatments as determined by Wilcoxon’s rank-sum test with Bonferroni’s correction (*P* < 0.05). There was no statistical difference among the treatments.

**Supplemental Figure 6.** Hyphal branching-inducing activity of other secoiridoid glucosides and geraniol. A, Box plots indicating the number of hyphal branches in *R. irregularis*. *R. irregularis* spores were treated with distilled water (Mock), 100 nM GR24, or the indicated concentrations of secoiridoid glucosides. Different letters indicate significant differences among treatments as determined by Wilcoxon’s rank-sum test with Bonferroni’s correction (*P* < 0.05, *n* = 12–19). B, Hyphal branching-inducing activity of 0.01% ethanol (Mock; cyan) and geraniol (yellow) in *R. irregularis*. There was no difference among treatments as determined by Wilcoxon’s rank-sum test with Bonferroni’s correction (*n* = 8–10). Data are shown as box plots with the 25th–75th percentiles (box), median (center line inside the box), and range (whiskers).

**Supplemental Figure 7.** Response of *R. clarus* and *G. margarita* to GPS and SWM. A, The number of hyphal branches of *R. clarus* treated with distilled water (Mock), 100 nM GR24, 1–100 nM GPS, or 1–100 nM SWM. Data are shown as box plots with the 25th–75th percentiles (box), median (center line inside the box), and range (whiskers). Different letters indicate significant differences among treatments as determined by Wilcoxon’s rank-sum test with Bonferroni’s correction (*P* < 0.05, *n* = 6–13). B, *G. margarita* hyphae treated with water, 0.1–10 µg/disk GPS, or SWM featured no hyphal branches. An aliquot (30 µL) of root exudates collected from *T. pratense* (red clover) was loaded onto a disk as a positive control. Arrows indicate the direction of primary hyphal elongation. Arrowheads denote newly formed hyphal branches after 24 h. Scale bars: 1 mm.

**Supplemental Figure 8.** HPLC analysis of methanol extracts collected from *L. japonicus* and chives. Methanol extracts of *L. japonicus roots* (orange line) and chives (blue line) showed no peaks that matched SWM (*Rt* 4.6 min) and GPS (*Rt* 5.7 min) standards (black line). Each extract was prepared at 50 mg root fresh weight mL^−1^.

**Supplemental Table S1.** Total DEGs in GA-treated *E. grandiflorum* mycorrhizal roots.

**Supplemental Table S2.** GO enrichment analysis of GA-treated *E. grandiflorum* mycorrhizal roots.

**Supplemental Table S3.** Expression pattern of secoiridoid pathway genes in *E. grandiflorum*.

**Supplemental Table S4.** Total DEGs in *R. irregularis* treated with GPS and GR24.

**Supplemental Table S5.** GO enrichment analysis on *R. irregularis* treated with GPS and GR24.

**Supplemental Table S6.** Results of trimming, mapping, and counting of RNA-seq reads.

## References

Akiyama K, Matsuzaki K, Hayashi H (2005) Plant sesquiterpenes induce hyphal branching in arbuscular mycorrhizal fungi. Nature 435: 824–827

Akiyama K, Ogasawara S, Ito S, Hayashi H (2010) Structural requirements of strigolactones for hyphal branching in AM fungi. Plant Cell Physiol 51: 1104–1117

Bakker PAHM, Pieterse CMJ, de Jonge R, Berendsen RL (2018) The soil-borne legacy. Cell 172: 1178–1180

Besserer A, Becard G, Jauneau A, Roux C, Sejalon-Delmas N (2008) GR24, a synthetic analog of strigolactones, stimulates the mitosis and growth of the arbuscular mycorrhizal fungus Gigaspora rosea by boosting its energy metabolism. Plant Physiol 148: 402–413

Besserer A, Puech-Pages V, Kiefer P, Gomez-Roldan V, Jauneau A, Roy S, Portais JC, Roux C, Becard G, Sejalon-Delmas N (2006) Strigolactones stimulate arbuscular mycorrhizal fungi by activating mitochondria. PLoS Biol 4: e226

Božunović J, Skorić M, Matekalo D, Živković S, Dragićević M, Aničić N, Filipović B, Banjanac T, Šiler B, Mišić D (2019) Secoiridoids metabolism response to wounding in common centaury (*Centaurium erythraea* Rafn) leaves. In Plants, Vol 8

Brundrett MC, Tedersoo L (2018) Evolutionary history of mycorrhizal symbioses and global host plant diversity. New Phytol 220: 1108–1115

Buchfink B, Reuter K, Drost H-G (2021) Sensitive protein alignments at tree-of-life scale using DIAMOND. Nat Methods 18: 366–368

Buwalda JG, Goh KM (1982) Host-fungus competition for carbon as a cause of growth depressions in vesicular-arbuscular mycorrhizal ryegrass. Soil Biol Biochem 14: 103–106

Cao X, Guo X, Yang X, Wang H, Hua W, He Y, Kang J, Wang Z (2016) Transcriptional responses and gentiopicroside biosynthesis in methyl jasmonate-treated *Gentiana macrophylla* seedlings. PLoS One 11: e0166493

Chen S, Zhou Y, Chen Y, Gu J (2018) fastp: an ultra-fast all-in-one FASTQ preprocessor. Bioinformatics 34: i884–i890

Cohen M, Prandi C, Occhiato EG, Tabasso S, Wininger S, Resnick N, Steinberger Y, Koltai H, Kapulnik Y (2013) Structure–function relations of strigolactone analogs: activity as plant hormones and plant interactions. Mol Plant 6: 141–152

Conesa A, Götz S, García-Gómez JM, Terol J, Talón M, Robles M (2005) Blast2GO: a universal tool for annotation, visualization and analysis in functional genomics research. Bioinformatics 21: 3674–3676

Cosme M, Fernández I, Declerck S, van der Heijden MGA, Pieterse CMJ (2021) A coumarin exudation pathway mitigates arbuscular mycorrhizal incompatibility in Arabidopsis thaliana. Plant Mol Biol 106: 319–334

Davière JM, Achard P (2013) Gibberellin signaling in plants. Development 140: 1147–1151

De Luca V, Salim V, Thamm A, Masada SA, Yu F (2014) Making iridoids/secoiridoids and monoterpenoid indole alkaloids: progress on pathway elucidation. Curr Opin Plant Biol 19: 35–42

de Saint Germain A, Bonhomme S, Boyer F-D, Rameau C (2013) Novel insights into strigolactone distribution and signalling. Curr Opin Plant Biol 16: 583–589

Dobin A, Davis CA, Schlesinger F, Drenkow J, Zaleski C, Jha S, Batut P, Chaisson M, Gingeras TR (2013) STAR: ultrafast universal RNA-seq aligner. Bioinformatics 29: 15–21

Dobler S, Petschenka G, Pankoke H (2011) Coping with toxic plant compounds - The insect’s perspective on iridoid glycosides and cardenolides. Phytochem 72: 1593–1604

Egusa M, Parada R, Aklog YF, Ifuku S, Kaminaka H (2019) Nanofibrillation enhances the protective effect of crab shells against Fusarium wilt disease in tomato. Int J Biol Macromol 128: 22–27

Foo E, Ross JJ, Jones WT, Reid JB (2013) Plant hormones in arbuscular mycorrhizal symbioses: an emerging role for gibberellins. Ann Bot 111: 769–779

Fujimatsu T, Endo K, Yazaki K, Sugiyama A (2020) Secretion dynamics of soyasaponins in soybean roots and effects to modify the bacterial composition. Plant Direct 4: e00259

Furlan V, Bartschi H, Fortin JA (1980) Media for density gradient extraction of endomycorrhizal spores. Trans Br Mycol Soc 75: 336–338

Ge S, He L, Jin L, Xia X, Li L, Ahammed GJ, Qi Z, Yu J, Zhou Y (2022) Light-dependent activation of HY5 promotes mycorrhizal symbiosis in tomato by systemically regulating strigolactone biosynthesis. New Phytol 233: 1900–1914

Genre A, Chabaud M, Faccio A, Barker DG, Bonfante P (2008) Prepenetration apparatus assembly precedes and predicts the colonization patterns of arbuscular mycorrhizal fungi within the root cortex of both Medicago truncatula and Daucus carota. Plant Cell 20: 1407–1420

Gutjahr C (2014) Phytohormone signaling in arbuscular mycorhiza development. Curr Opin Plant Biol 20: 26–34

Hildebrandt U, Janetta K, Bothe H (2002) Towards growth of arbuscular mycorrhizal fungi independent of a plant host. Appl Environ Microbiol 68: 1919–1924

Ito S, Yamagami D, Umehara M, Hanada A, Yoshida S, Sasaki Y, Yajima S, Kyozuka J, Ueguchi-Tanaka M, Matsuoka M, et al. (2017) Regulation of strigolactone biosynthesis by gibberellin signaling. Plant Physiol 174: 1250–1259

Kameoka H, Maeda T, Okuma N, Kawaguchi M (2019) Structure-specific regulation of nutrient transport and metabolism in arbuscular mycorrhizal fungi. Plant Cell Physiol 60: 2272–2281

Kameoka H, Tsutsui I, Saito K, Kikuchi Y, Handa Y, Ezawa T, Hayashi H, Kawaguchi M, Akiyama K (2019) Stimulation of asymbiotic sporulation in arbuscular mycorrhizal fungi by fatty acids. Nat Microbiol 4: 1654–1660

Kaur S, Campbell BJ, Suseela V (2022) Root metabolome of plant–arbuscular mycorrhizal symbiosis mirrors the mutualistic or parasitic mycorrhizal phenotype. New Phytol 234: 672–687

Ketudat Cairns JR, Esen A (2010) β-Glucosidases. Cell Mol Life Sci 67: 3389–3405

Kim DH, Kim BR, Kim JY, Jeong YC (2000) Mechanism of covalent adduct formation of aucubin to proteins. Toxicol Let 114: 181–188

Kobae Y, Kameoka H, Sugimura Y, Saito K, Ohtomo R, Fujiwara T, Kyozuka J (2018) Strigolactone biosynthesis genes of rice are required for the punctual entry of arbuscular mycorrhizal fungi into the roots. Plant Cell Physiol 59: 544–553

Kobayashi Y, Maeda T, Yamaguchi K, Kameoka H, Tanaka S, Ezawa T, Shigenobu S, Kawaguchi M (2018) The genome of *Rhizophagus clarus* HR1 reveals a common genetic basis for auxotrophy among arbuscular mycorrhizal fungi. BMC Genom 19: 465

Kodama K, Rich MK, Yoda A, Shimazaki S, Xie X, Akiyama K, Mizuno Y, Komatsu A, Luo Y, Suzuki H, et al. (2022) An ancestral function of strigolactones as symbiotic rhizosphere signals. Nat Commun 13: 3974

Konno K, Hirayama C, Yasui H, Nakamura M (1999) Enzymatic activation of oleuropein: a protein crosslinker used as a chemical defense in the privet tree. Proc Natl Acad Sci 96: 9159–9164

Kretzschmar T, Kohlen W, Sasse J, Borghi L, Schlegel M, Bachelier JB, Reinhardt D, Bours R, Bouwmeester HJ, Martinoia E (2012) A petunia ABC protein controls strigolactone-dependent symbiotic signalling and branching. Nature 483: 341–344

Kumarasamy Y, Nahar L, Sarker SD (2003) Bioactivity of gentiopicroside from the aerial parts of *Centaurium erythraea*. Fitoterapia 74: 151–154

Lendenmann M, Thonar C, Barnard RL, Salmon Y, Werner RA, Frossard E, Jansa J (2011) Symbiont identity matters: carbon and phosphorus fluxes between *Medicago truncatula* and different arbuscular mycorrhizal fungi. Mycorrhiza 21: 689–702

Li T, Yu X, Ren Y, Kang M, Yang W, Feng L, Hu Q (2022) The chromosome-level genome assembly of *Gentiana dahurica* (Gentianaceae) provides insights into gentiopicroside biosynthesis. DNA Res 29: dsac008

Liao Y, Smyth GK, Shi W (2014) featureCounts: an efficient general purpose program for assigning sequence reads to genomic features. Bioinformatics 30: 923–930

Lichius A, Berepiki A, Read ND (2011) Form follows function – The versatile fungal cytoskeleton. Fungal Biol 115: 518–540

Luginbuehl LH, Oldroyd GED (2017) Understanding the arbuscule at the heart of endomycorrhizal symbioses in plants. Curr Biol 27: R952–R963

Maeda T, Kobayashi Y, Kameoka H, Okuma N, Takeda N, Yamaguchi K, Bino T, Shigenobu S, Kawaguchi M (2018) Evidence of non-tandemly repeated rDNAs and their intragenomic heterogeneity in Rhizophagus irregularis. Commun Biol 1: 87

Nadal M, Sawers R, Naseem S, Bassin B, Kulicke C, Sharman A, An G, An K, Ahern KR, Romag A, et al. (2017) An N-acetylglucosamine transporter required for arbuscular mycorrhizal symbioses in rice and maize. Nat Plants 3: 17073

Nagata M, Yamamoto N, Shigeyama T, Terasawa Y, Anai T, Sakai T, Inada S, Arima S, Hashiguchi M, Akashi R, et al. (2015) Red/far red light controls arbuscular mycorrhizal colonization via jasmonic acid and strigolactone signaling. Plant Cell Physiol 56: 2100–2109

Nakabayashi R, Takeda-Kamiya N, Yamada Y, Mori T, Uzaki M, Nirasawa T, Toyooka K, Saito K (2021) A multimodal metabolomics approach using imaging mass spectrometry and liquid chromatography-tandem mass spectrometry for spatially characterizing monoterpene indole alkaloids secreted from roots. Plant Biotechnol 38: 305–310

Nakayasu M, Ohno K, Takamatsu K, Aoki Y, Yamazaki S, Takase H, Shoji T, Yazaki K, Sugiyama A (2021) Tomato roots secrete tomatine to modulate the bacterial assemblage of the rhizosphere. Plant Physiol 186: 270–284

Nouri E, Surve R, Bapaume L, Stumpe M, Chen M, Zhang Y, Ruyter-Spira C, Bouwmeester H, Glauser G, Bruisson S, et al. (2021) Phosphate suppression of arbuscular mycorrhizal symbiosis involves gibberellic acid signalling. Plant Cell Physiol 62: 959–970

Pimprikar P, Carbonnel S, Paries M, Katzer K, Klingl V, Bohmer MJ, Karl L, Floss DS, Harrison MJ, Parniske M, et al. (2016) A CCaMK-CYCLOPS-DELLA complex activates transcription of *RAM1* to regulate arbuscule branching. Curr Biol 26: 987–998

Rai A, Hirakawa H, Nakabayashi R, Kikuchi S, Hayashi K, Rai M, Tsugawa H, Nakaya T, Mori T, Nagasaki H, et al. (2021) Chromosome-level genome assembly of Ophiorrhiza pumila reveals the evolution of camptothecin biosynthesis. Nat Commun 12: 405

Rai A, Nakamura M, Takahashi H, Suzuki H, Saito K, Yamazaki M (2016) High-throughput sequencing and de novo transcriptome assembly of *Swertia japonica* to identify genes involved in the biosynthesis of therapeutic metabolites. Plant Cell Rep 35: 2091–2111

Robinson MD, McCarthy DJ, Smyth GK (2010) edgeR: a Bioconductor package for differential expression analysis of digital gene expression data. Bioinformatics 26: 139–140

Shirasawa K, Arimoto R, Hirakawa H, Ishimori M, Ghelfi A, Miyasaka M, Endo M, Kawabata S, Isobe SN (2023) Chromosome-scale genome assembly of *Eustoma grandiflorum*, the first complete genome sequence in the genus *Eustoma*. G3 Genes|Genomes|Genetics 13: jkac329

Šiler B, Mišić D, Nestorović J, Banjanac T, Glamočlija J, Soković M, Ćirić A (2010) Antibacterial and antifungal screening of *Centaurium pulchellum* crude extracts and main secoiridoid compounds. Nat Prod Commun 5: 1934578X1000501001

Soler-Rivas C, Espín JC, Wichers HJ (2000) Oleuropein and related compounds. J Sci Food Agric 80: 1013–1023

Takeda N, Handa Y, Tsuzuki S, Kojima M, Sakakibara H, Kawaguchi M (2015) Gibberellins interfere with symbiosis signaling and gene expression and alter colonization by arbuscular mycorrhizal fungi in *Lotus japonicus*. Plant Physiol 167: 545–557

Tisserant E, Malbreil M, Kuo A, Kohler A, Symeonidi A, Balestrini R, Charron P, Duensing N, Frei dit Frey N, Gianinazzi-Pearson V, et al. (2013) Genome of an arbuscular mycorrhizal fungus provides insight into the oldest plant symbiosis. Proc Natl Acad Sci 110: 20117–20122

Tominaga T, Miura C, Sumigawa Y, Hirose Y, Yamaguchi K, Shigenobu S, Mine A, Kaminaka H (2021) Conservation and diversity in gibberellin-mediated transcriptional responses among host plants forming distinct arbuscular mycorrhizal morphotypes. Front Plant Sci 12: 795695

Tominaga T, Miura C, Takeda N, Kanno Y, Takemura Y, Seo M, Yamato M, Kaminaka H (2020) Gibberellin promotes fungal entry and colonization during *Paris*-type arbuscular mycorrhizal symbiosis in *Eustoma grandiflorum*. Plant Cell Physiol 61: 565–575

Tominaga T, Yamaguchi K, Shigenobu S, Yamato M, Kaminaka H (2020) The effects of gibberellin on the expression of symbiosis-related genes in *Paris*-type arbuscular mycorrhizal symbiosis in *Eustoma grandiflorum*. Plant Signal Behav 15: 1784544

Tsuzuki S, Handa Y, Takeda N, Kawaguchi M (2016) Strigolactone-induced putative secreted protein 1 Is required for the establishment of symbiosis by the arbuscular mycorrhizal fungus *Rhizophagus irregularis*. Mol Plant Microbe Interact 29: 277–286

Ueno K, Furumoto T, Umeda S, Mizutani M, Takikawa H, Batchvarova R, Sugimoto Y (2014) Heliolactone, a non-sesquiterpene lactone germination stimulant for root parasitic weeds from sunflower. Phytochem 108: 122–128

Wang S-C, Tseng T-Y, Huang C-M, Tsai T-H (2004) Gardenia herbal active constituents: applicable separation procedures. J Chromatogr B 812: 193–202

Yang C, Li L (2017) Hormonal regulation in shade avoidance. Front Plant Sci 8: 1527

Yoneyama K, Xie X, Kusumoto D, Sekimoto H, Sugimoto Y, Takeuchi Y, Yoneyama K (2007) Nitrogen deficiency as well as phosphorus deficiency in sorghum promotes the production and exudation of 5-deoxystrigol, the host recognition signal for arbuscular mycorrhizal fungi and root parasites. Planta 227: 125–132

Yoneyama K, Xie X, Yoneyama K, Kisugi T, Nomura T, Nakatani Y, Akiyama K, McErlean CSP (2018) Which are the major players, canonical or non-canonical strigolactones? J Exp Bot 69: 2231–2239

Yu F, Yu F, Li R, Wang R (2004) Inhibitory effects of the *Gentiana macrophylla* (Gentianaceae) extract on rheumatoid arthritis of rats. J Ethnopharmacol 95: 77–81

Zhong Y, Xun W, Wang X, Tian S, Zhang Y, Li D, Zhou Y, Qin Y, Zhang B, Zhao G, et al. (2022) Root-secreted bitter triterpene modulates the rhizosphere microbiota to improve plant fitness. Nat Plants 8: 887–896

Zhou T, Bai G, Hu Y, Ruhsam M, Yang Y, Zhao Y (2022) *De novo* genome assembly of the medicinal plant *Gentiana macrophylla* provides insights into the genomic evolution and biosynthesis of iridoids. DNA Res 29: dsac034

