## Supplemental Figure 1-8 for "Monoterpene glucosides accumulated in *Eustoma grandiflorum* roots promote hyphal branching in arbuscular mycorrhizal fungi"

### Fig. S1

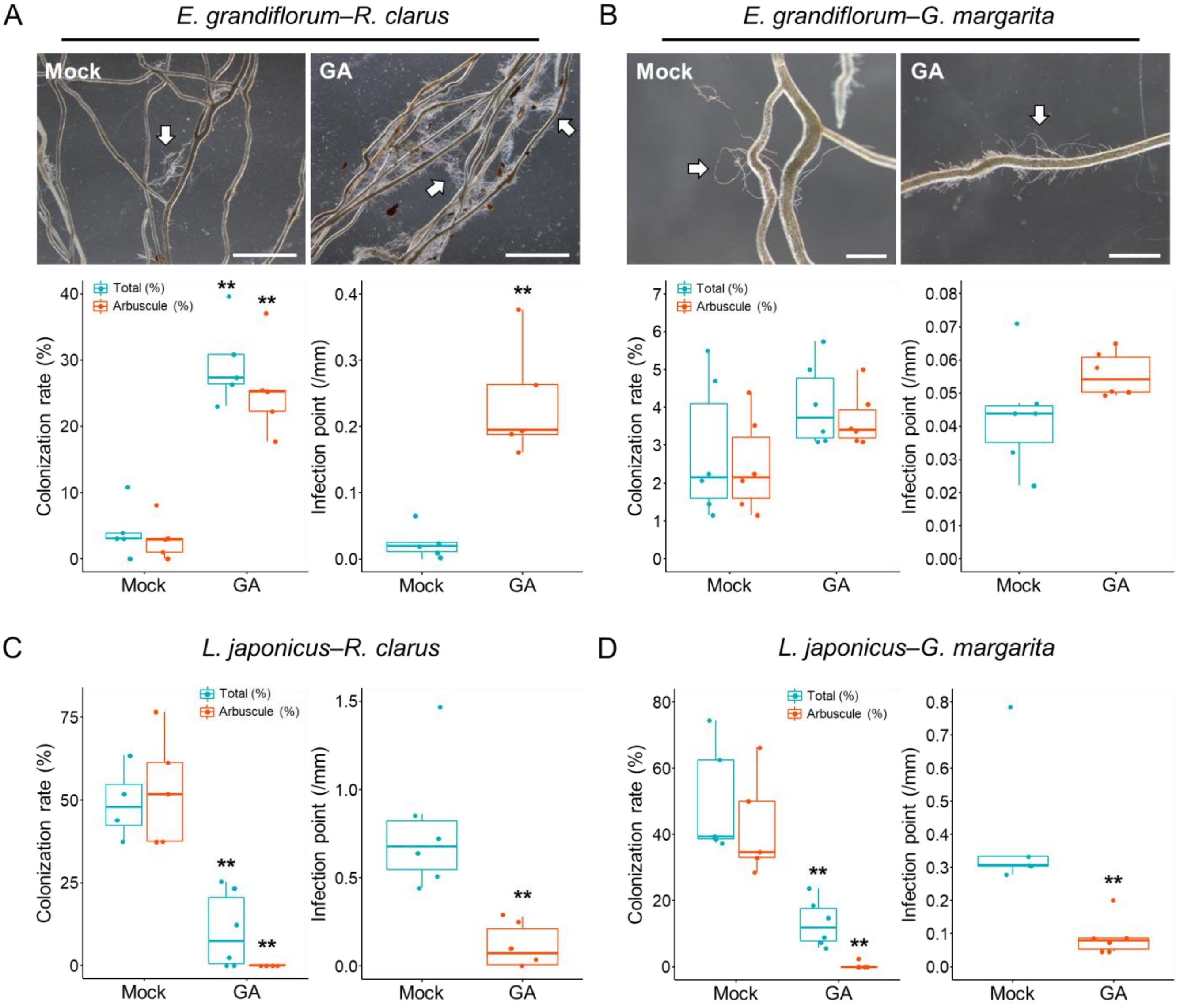


**Supplemental Figure S1.** Different effects of GA treatment on AM symbioses between *E. grandiflorum* and *L. japonicus*. *E. grandiflorum* and *L. japonicus* roots inoculated with *R. clarus* or *G. margarita* were observed at 5 weeks post-inoculation. The plants were treated with 0.01% ethanol (Mock) or 1 μM $GA_3$ (GA). A and B, *E. grandiflorum* roots were colonized by *R. clarus* (A) and *G. margarita* (B). Arrows denote extraradical hyphae adhering to *E. grandiflorum* roots. Scale bars, 5 mm (A) and 1 mm (B). Below graphs show the colonization rates (%) and hyphopodia number ($mm^{-1}$) of *R. clarus* (A) and *G. margarita* (B) infecting *E. grandiflorum*. C and D, Colonization rates (%) and hyphopodia number ($mm^{-1}$) of *R. clarus* (C) and *G. margarita* (D) infecting *L. japonicus*. Green and orange plots present the total hyphal colonization and arbuscule formation rates, respectively. Data are shown as box plots with the 25th –75th percentiles (box), median (center line inside the box), and range (whiskers). Asterisks indicate significant differences in GA-treated plants compared to mock-treated plant as determined using Wilcoxon's rank-sum test (**: $P < 0.01$, $n = 5–6$).

### Fig. S2

A

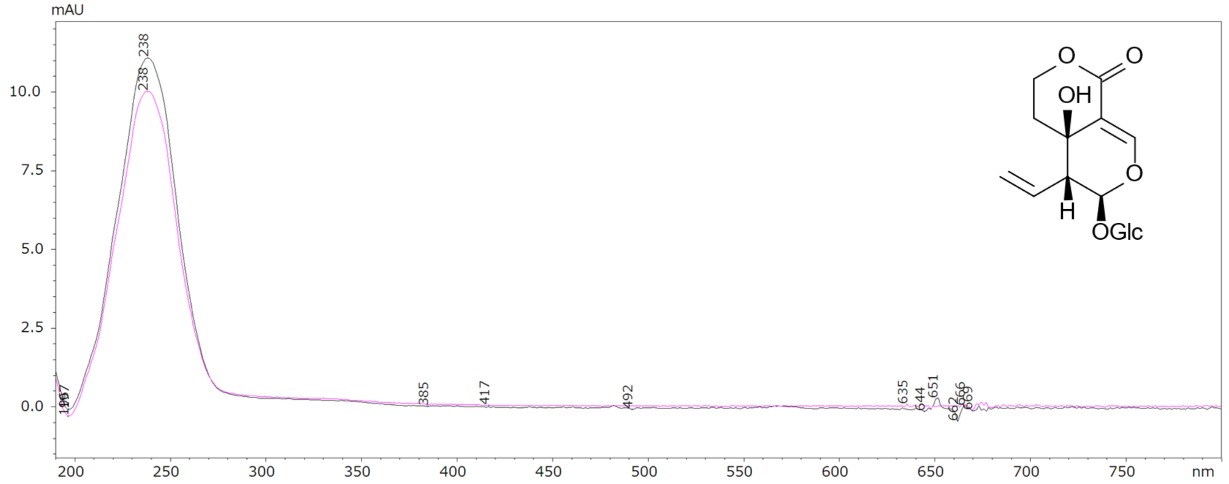


B

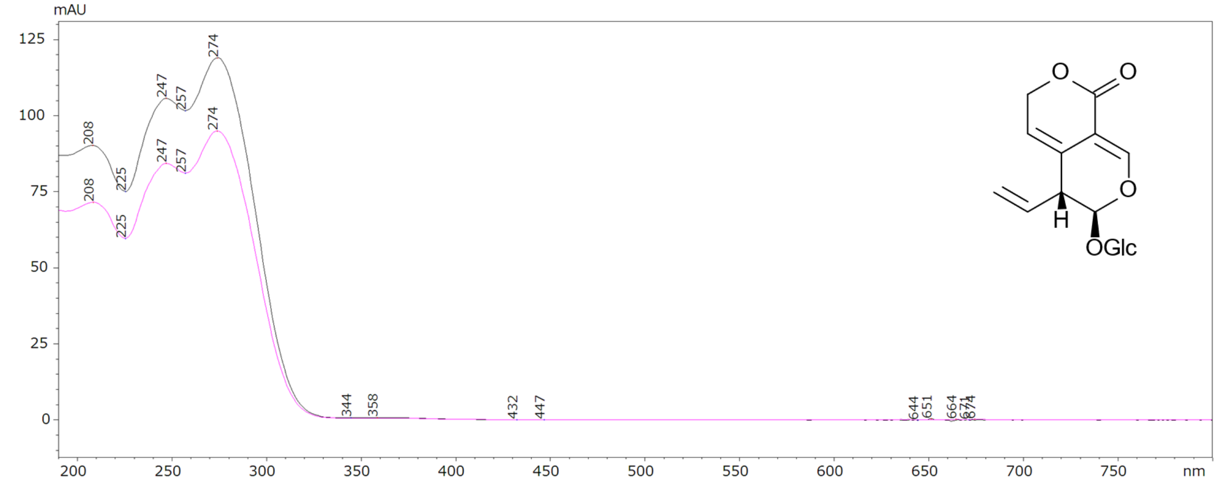


**Supplemental Figure 2.** HPLC analyses of peaks 1 and 2 in Fig. 1C. A and B, Magenta lines represent the UV spectra of peaks 1 (A) and 2 (B) from the methanol extracts of 6-week-old axenic *E. grandiflorum* roots (Fig. 1C). Black lines indicate the UV spectra of SW (A) and GPS (B) standards. UV spectra of peaks 1 and 2 matched the SWM and GPS standard spectra (99.9% and 100%), respectively.

Fig. S3

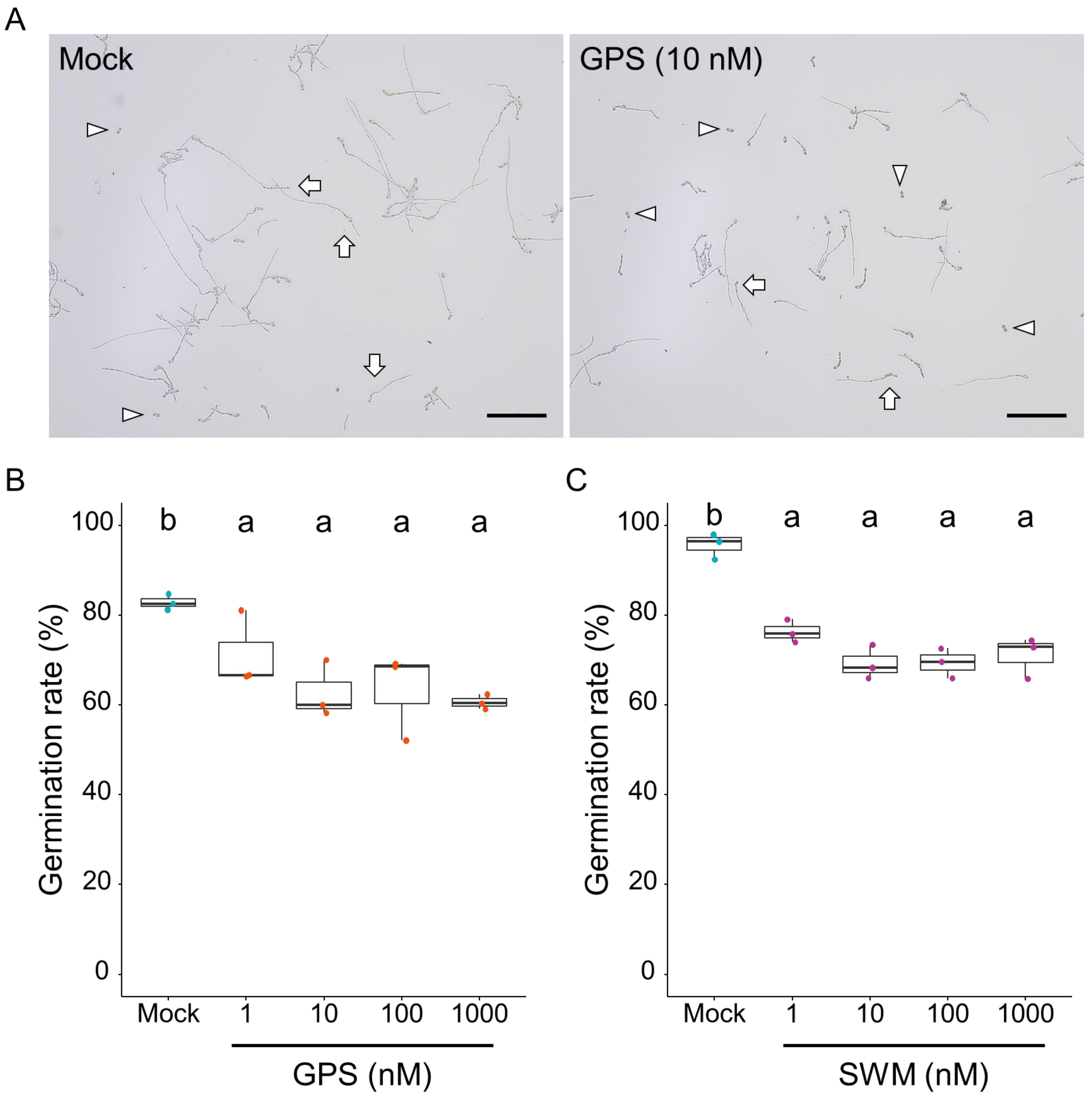


**Supplemental Figure 3.** Antifungal activity of GPS and SWM against *F. oxysporum*. A, Images of *F. oxysporum* f. sp. *lycopersici* bud cells treated with distilled water (left) or 10 nM GPS (right). Arrows indicate germinating bud cells, and arrowheads denote bud cells that did not germinate by 12 h. Scale bars, 100 µm. B and C, Germination rates of *F. oxysporum* bud cells treated with 1–1000 nM GPS (B) or SWM (C). Data are shown as box plots with the 25th–75th percentiles (box), the (center line inside the box), and range (whiskers). Different letters indicate significant differences among treatments as determined using the Tukey test ($P < 0.001$, $n = 3$).

Fig. S4

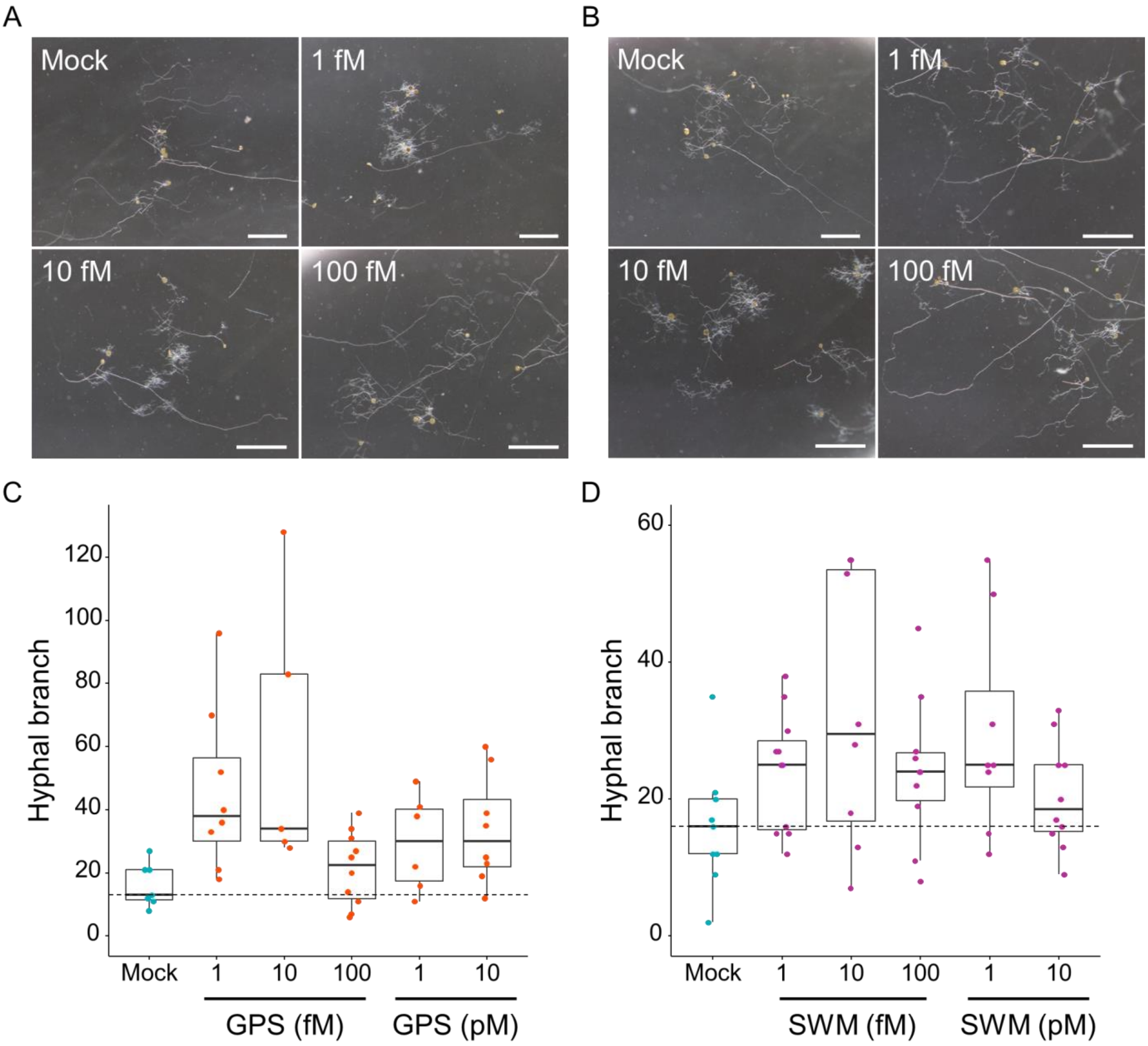


**Supplemental Figure 4.** Hyphal branching induced by GPS and SWM in the femtomolar range. A and B, Images showing *R. irregularis* hyphae treated with femtomolar level GPS (A) and SWM (B). Distilled water was used as a mock control. Scale bars, 1 mm. C and D, The number of hyphal branches of *R. irregularis* treated with GPS (C) and SWM (D). There was no difference among treatments as determined using Wilcoxon's rank-sum test with Bonferroni's correction ($n$ = 5–11). Data are shown as box plots with the 25th–75th percentiles (box), median (center line inside the box), and range (whiskers).

Fig. S5

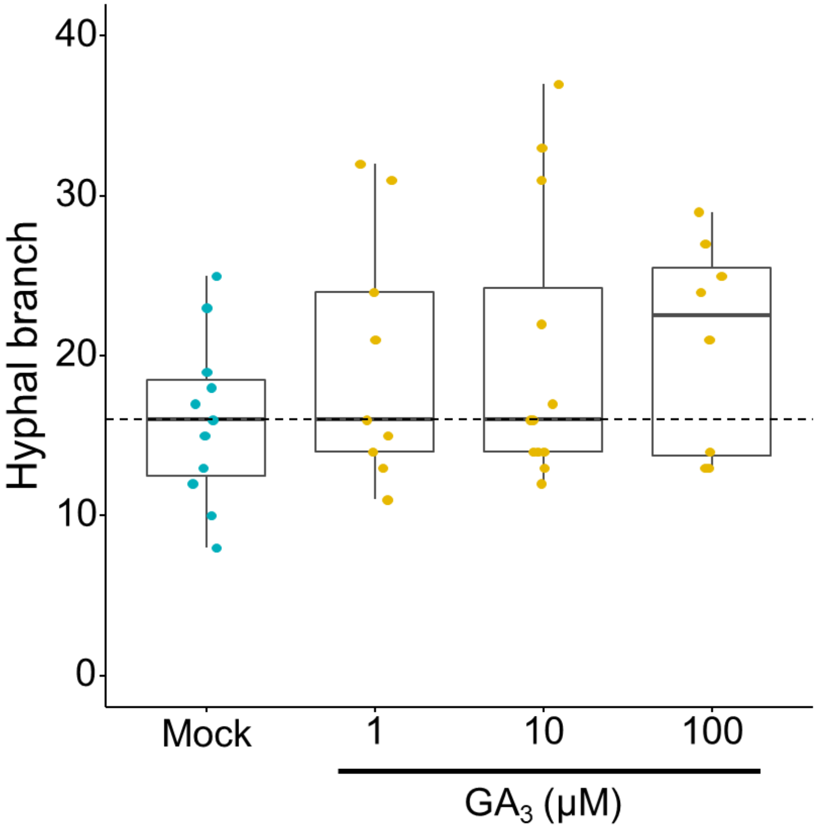


**Supplemental Figure 5.** Hyphal branching-inducing activity of GA. The effect of exogenous $GA_3$ on *R. irregularis* hyphal branching. For the mock treatment, 0.01% ethanol was supplied to the fungus ($n$ = 9–13). Different letters indicate significant differences among treatments as determined by Wilcoxon’s rank-sum test with Bonferroni’s correction ($P$ < 0.05). There was no statistical difference among the treatments.

### Fig. S6

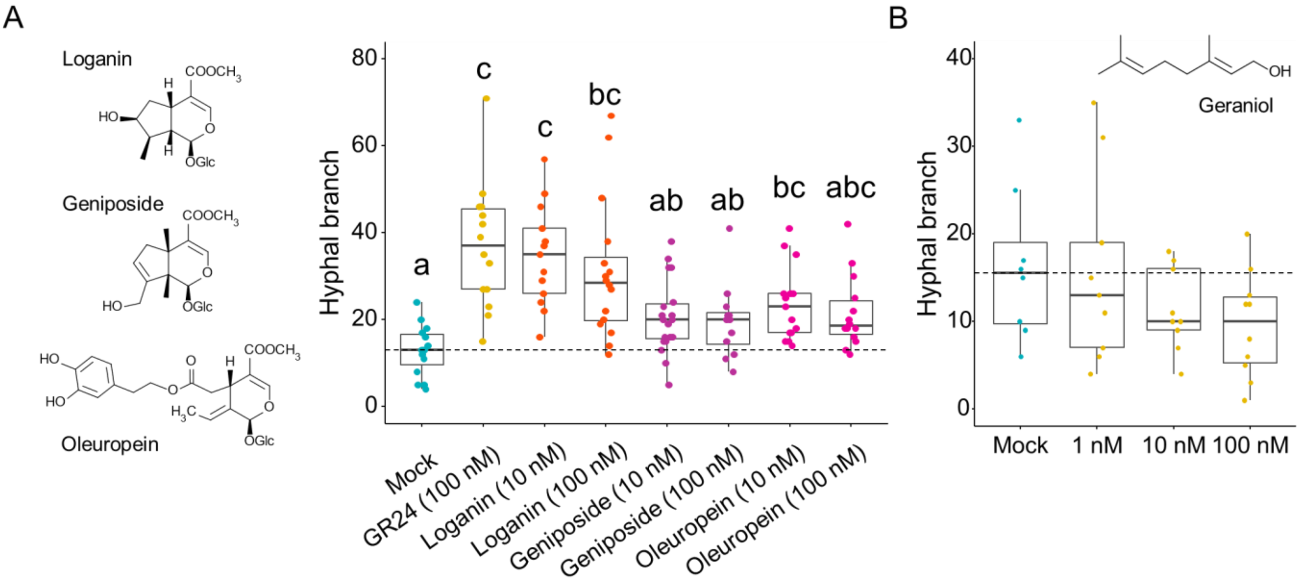


**Supplemental Figure 6.** Hyphal branching-inducing activity of other secoiridoid glucosides and geraniol. A, Box plots indicating the number of hyphal branches in *R. irregularis*. *R. irregularis* spores were treated with distilled water (Mock), 100 nM GR24, or the indicated concentrations of secoiridoid glucosides. Different letters indicate significant differences among treatments as determined by Wilcoxon's rank-sum test with Bonferroni's correction ($P < 0.05$, $n = 12–19$). B, Hyphal branching-inducing activity of 0.01% ethanol (Mock; cyan) and geraniol (yellow) in *R. irregularis*. There was no difference among treatments as determined by Wilcoxon's rank-sum test with Bonferroni's correction ($n = 8–10$). Data are shown as box plots with the 25th–75th percentiles (box), median (center line inside the box), and range (whiskers).

Fig. S7

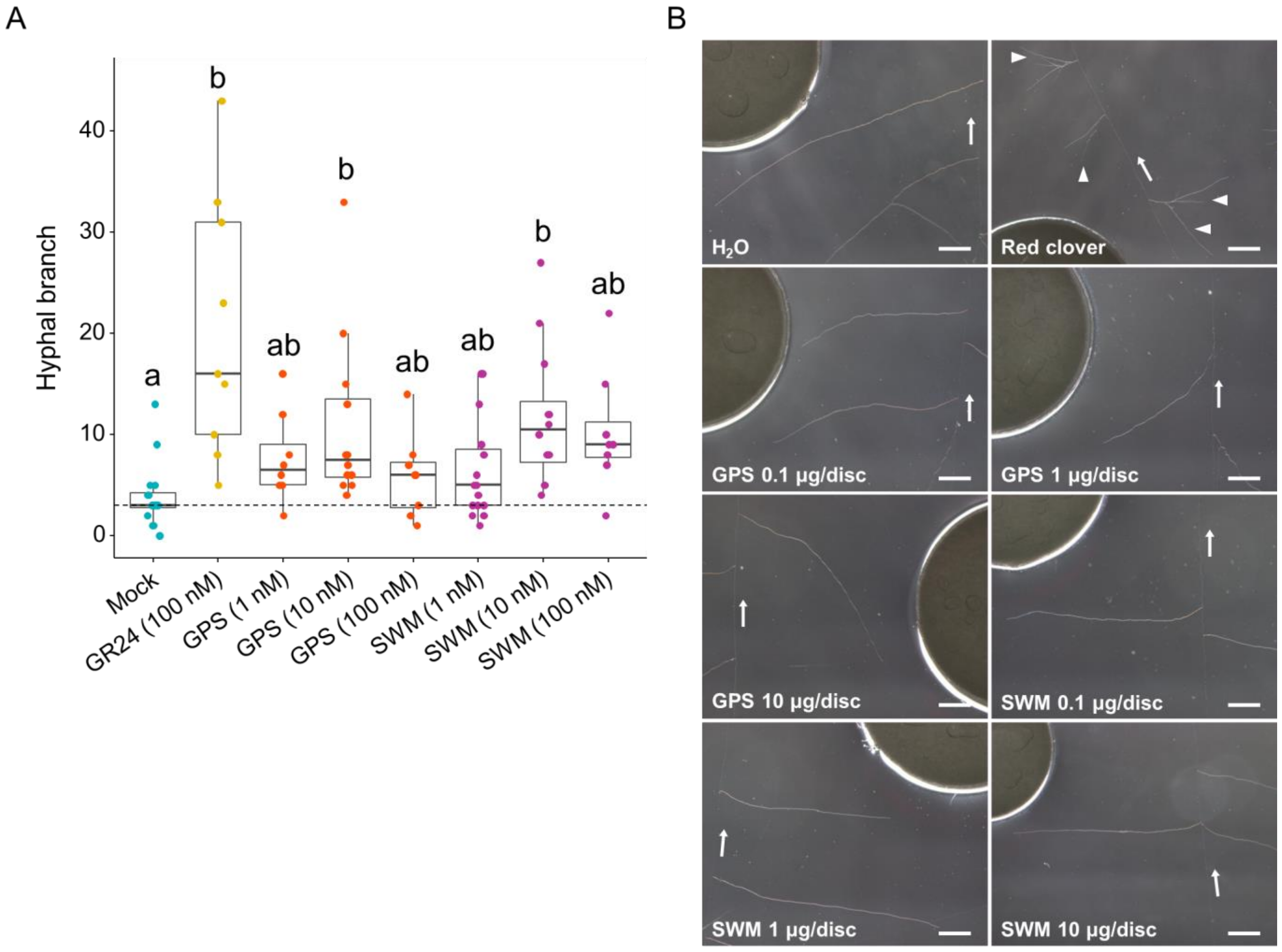


**Supplemental Figure 7.** Response of *R. clarus* and *G. margarita* to GPS and SWM. A, The number of hyphal branches of *R. clarus* treated with distilled water (Mock), 100 nM GR24, 1–100 nM GPS, or 1–100 nM SWM. Data are shown as box plots with the 25th–75th percentiles (box), median (center line inside the box), and range (whiskers). Different letters indicate significant differences among treatments as determined by Wilcoxon's rank-sum test with Bonferroni's correction ($P < 0.05$, $n = 6–13$). B, *G. margarita* hyphae treated with water, 0.1–10 µg/disk GPS, or SWM featured no hyphal branches. An aliquot (30 µL) of root exudates collected from *T. pratense* (red clover) was loaded onto a disk as a positive control. Arrows indicate the direction of primary hyphal elongation. Arrowheads denote newly formed hyphal branches after 24 h. Scale bars: 1 mm.

Fig. S8

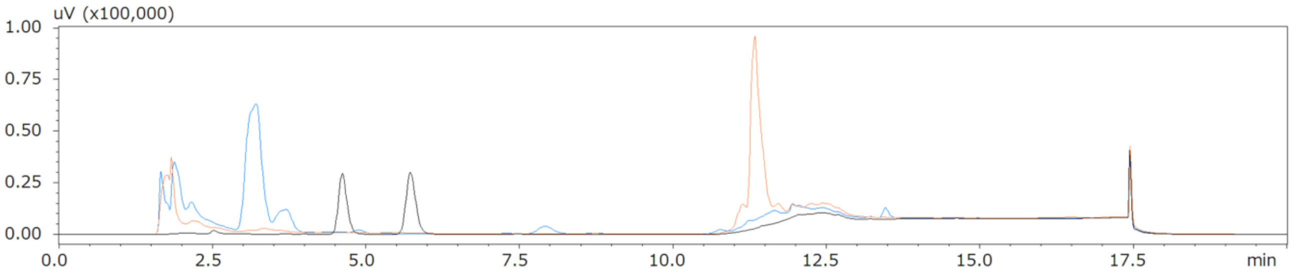


**Supplemental Figure 8.** HPLC analysis of methanol extracts collected from *L. japonicus* and chives. Methanol extracts of *L. japonicus roots* (orange line) and chives (blue line) showed no peaks that matched SWM (*Rt* 4.6 min) and GPS (*Rt* 5.7 min) standards (black line). Each extract was prepared at 50 mg root fresh weight $mL^{-1}$.
